# Hidden Diversity in *Enterococcus faecalis* Revealed by CRISPR2 Screening: Eco-evolutionary Insights into a Novel Subspecies

**DOI:** 10.1101/2025.05.05.652174

**Authors:** Vitor Luis Macena Leite, Adriana Rocha Faria, Clara Ferreira Guerra, Stephanie da Silva Rodrigues Souza, Andréa de Andrade Rangel Freitas, Jaqueline Martins Morais, Vânia Lúcia Carreira Merquior, Paul J. Planet, Lúcia Martins Teixeira

## Abstract

*Enterococcus faecalis* is a commensal bacterium that colonizes the gut of humans and animals, and a major opportunistic pathogen, known for causing multidrug-resistant healthcare-associated infections (HAIs). Its ability to thrive in diverse environments and disseminate antimicrobial resistance genes (ARGs) across ecological niches highlights the importance of understanding its ecological, evolutionary, and epidemiological dynamics. The CRISPR2 locus has been used as a valuable marker for assessing clonality and phylogenetic relationships in *E. faecalis*. In this study, we identified a group of *E. faecalis* strains lacking CRISPR2, forming a distinct, well-supported clade. We demonstrate that this clade meets the genomic criteria for classification as a novel subspecies, here referred to as “subspecies B”. Through a comprehensive pangenome analysis and comparative genomics, we explored the adaptive ecological traits underlying this diversification process, identifying clade-specific features and their predicted functional roles. Our findings suggest that the frequent isolation of subspecies B from meat products and processing facilities may reflect dissemination routes involving environmental contamination (e.g., water, plants, soil) from avian species. The absence of key virulence traits required for pathogenicity in mammals, particularly in humans, and the lack of clinically relevant resistance determinants indicate that subspecies B may currently pose minimal threat to public health compared to the broadly disseminated “subspecies A”. Nevertheless, the unclear potential for genetic exchange between these subspecies, and the frequent association of subspecies B with food sources, calls for continued genomic surveillance of *E. faecalis* from a One Health perspective to detect and mitigate the emergence of high-risk variants in advance.

## INTRODUCTION

*Enterococcus faecalis* is a Gram-positive commensal bacterium commonly found colonizing the gastrointestinal tract (GIT) of humans and other animals (1). Its broad distribution, extending to soil, water, and other natural sources (2), is attributed to its exceptional tolerance to adverse environmental conditions, including variations in temperature, pH, and exposure to antimicrobials (3, 4). As an opportunistic pathogen, *E. faecalis* is a major cause of healthcare-associated infections (HAIs) globally, mainly due to its remarkable ability to acquire multiple antimicrobial resistance genes (ARGs) through horizontal gene transfer (HGT), hampering treatment outcomes (5, 6).

A key factor in the genomic plasticity of *E. faecalis* is its ability to accumulate mobile genetic elements (MGEs), a process counterbalanced by Clustered Regularly Interspaced Short Palindromic Repeats (CRISPR) and their associated Cas proteins (CRISPR-Cas) (5). The CRISPR-Cas system functions as a bacterial adaptive immune mechanism, protecting against bacteriophage infections and limiting the uptake of exogenous DNA, including plasmids and other MGEs (7). The system’s activity relies on two core components: (i) the CRISPR array, composed of alternating direct repeats and short spacer sequences derived from foreign DNA, and (ii) Cas proteins, which mediate the recognition and cleavage of invading genetic material (7).

Three CRISPR loci have been described in *E. faecalis*: CRISPR1-Cas and CRISPR3-Cas, both predicted to be functional, and CRISPR2, an orphan locus lacking associated *cas* genes, rendering it incapable of defense against foreign DNA (5). Despite this, CRISPR2 has been widely regarded as a conserved feature of the *E. faecalis* genome, found in most isolates and considered part of the species’ core genome (8). This raises intriguing questions about its evolutionary maintenance, particularly given to evidence suggesting a potential regulatory role for CRISPR2 as a noncoding RNA (9). Indeed, transcriptomic analyses have confirmed CRISPR2 transcription, though its precise function remains unconfirmed (10, 11).

Previous studies have explored the discriminatory power of CRISPR2 arrays to assess *E. faecalis* genetic diversity (12–14). Given the system’s ubiquity and the variability of spacer sequences, CRISPR2 has been proposed as a useful marker for strain typing, especially in studies with limited resources where more comprehensive methods like whole-genome sequencing are not feasible. Notably, Hullahalli and colleagues (2015) demonstrated the value of CRISPR2 typing in providing additional phylogenetic resolution when coupled with multilocus sequence typing (MLST). However, while CRISPR2-based methods have proven valuable for investigating population diversity, studies have not yet explored potential associations between unique CRISPR2 signatures and ecological variation within closely related strains, especially in ubiquitous *E. faecalis* sequence-types (STs), such as STs 16, 21, and 40 (14, 15).

Given the generalist nature of *E. faecalis* and its exposure to diverse ecological pressures beyond healthcare settings, it is crucial to explore its population structure through molecular epidemiology approaches. Studies comprising isolates from a range of environments, host species, and geographical regions will deepen our understanding of the ecological and evolutionary dynamics shaping *E. faecalis* diversity (16). Thus, it is extremely worthy to explore inexpensive and feasible approaches providing complementary resolution to conventional typing methods (e.g., MLST), enabling the discrimination of subpopulations in early stages of ecological differentiation, which tend to be indistinguishable by multilocus sequence analyses targeting housekeeping genes (17, 18).

Initially, this study aimed to explore CRISPR2 sequence variability across a comprehensive dataset of *E. faecalis* genomes representing multiple STs and diverse isolation sources. However, an unexpected finding shifted the focus of our research. A subset of isolates from genetically related STs was found to completely lack CRISPR2. This prompted us to reframe the study with two main objectives. First, we aimed to validate whether this CRISPR2-negative group constitutes a cohesive unit of bacterial diversity, distinct both genetically and ecologically, while also clarifying its taxonomic and phylogenetic position in relation to well-characterized *E. faecalis* lineages. Second, through comparative pangenomic analysis, we sought to identify the differential genetic content between both groups and discuss potential selective pressures driving cladogenesis within *E. faecalis*. This genome-based approach offers novel insights into the complexity of *E. faecalis* population structure and niche breadth, identifying key areas for future studies on the ecological and evolutionary dynamics underlying genome divergence in this species.

## MATERIALS AND METHODS

### *Enterococcus faecalis* GENOMIC DATA

Initially, this study included 71 whole genome sequences of *E. faecalis* strains from our local bacterial culture collection, selected based on data availability rather than specific inclusion criteria. These strains were originally isolated from diverse sources, primarily in the state of Rio de Janeiro, Brazil, including 60 from hospitalized patients, 8 from coastal waters, and 3 from wild birds admitted to wildlife rehabilitation centers, with isolation dates ranging from 2005 to 2017. At this preliminary stage, the goal was to explore CRISPR2 sequence variability across a broad representation of *E. faecalis* genomes, without pre-established hypotheses regarding the ecological or clinical origins of the isolates.

Genomic DNA extraction, purification, and sequencing followed a previously described protocol (19), using the Illumina HiSeq 2500 platform (Illumina Inc., San Diego, CA, USA). Information on local bacterial strains and access details for their corresponding raw sequencing data are provided in the **S1 Dataset**. *De novo* genome assemblies were performed with the Unicycler v0.5.0 pipeline (20) on the BV-BRC suite (21), with default parameters and automatic read trimming by Trim Galore v0.6.5 (22).

In addition, 1,464 complete and draft *E. faecalis* genomes, available in GenBank until April 1, 2020, were downloaded for further analysis. All genomes were assigned sequence types (STs) using *in silico* MLST based on PubMLST typing schemes (23, 24). Targeted searches for genomes representing specific STs, flagged as relevant based on initial findings, were conducted in the updated GenBank database (up to May 30, 2022), resulting in the inclusion of three additional *E. faecalis* genomes isolated between 2015 and 2019. Throughout this study, specific subsets of genomes were analyzed to address emerging research questions as new patterns were identified. The rationale for these subsets is detailed alongside the corresponding results to provide contextual clarity and maintain a logical flow of information.

All *E. faecalis* genomes analyzed in this study, including strain metadata and accession numbers are shown in **S1 Dataset**. Genome annotation was carried out using Prokka v1.14.5 (25).

### CRISPR SCREENING

All *E. faecalis* genomes were screened for the presence of CRISPR2 loci using two complementary approaches: the CRISPRCasFinder online tool (26) and an *in silico* PCR program described in reference 27, employing previously described primer sets (28). CRISPRCasFinder detects candidate CRISPR-*cas* systems within a query genome, allowing to distinguish CRISPR2 loci from other *E. faecalis* CRISPR systems (CRISPR1-*cas* and CRISPR3-*cas*) using two criteria: (i) the presence of signature direct repeats (28); and ii) the absence of adjacent *cas* genes. Functional CRISPR-*cas* loci (CRISPR1-*cas* and CRISPR3-*cas*) identified by CRISPRCasFinder were also noted for subsequent analysis.

For CRISPR2 detection via *in silico* PCR, any amplification product was considered indicative of the locus’s presence, regardless of size, as the primers were designed to target its conserved flanking regions in the chromosome (28). This differs from the detection of CRISPR1-*cas* and CRISPR3-*cas* systems, which relies on primers targeting the *cas* genes. The variation in product size across strains reflects the number of spacers within the CRISPR2 array. CRISPR2 absence was confirmed only when both screening methods (CRISPRCasFinder and *in silico* PCR) failed to detect the array.

### DETECTION OF ANTIMICROBIAL RESISTANCE AND VIRULENCE-ASSOCIATED GENES

Genomes of interest were screened for ARGs and virulence genes against the Comprehensive Antibiotic Resistance Database (CARD; http://arpcard.mcmaster.ca) (29) and the Virulence Factor Database (VFDB; http://www.mgc.ac.cn/VFs/) (30), respectively, using ABRicate v1.0.0 (31) with default parameters.

### GENOME-BASED TAXONOMY

Taxonomic confirmation of the genomes of interest was conducted using the Type (Strain) Genome Server (TYGS), a high-throughput platform designed for genome-based taxonomic analysis (32, 33). This platform is interconnected with the List of Prokaryotic Names with Standing in Nomenclature (LPSN) database (34). The algorithm assigns each query genome to its corresponding species and subspecies based on established digital DNA:DNA hybridization (dDDH) thresholds — 70% for species delineation and 70-80% for subspecies — (35, 36), using pairwise comparisons against the closest type strains in the TYGS database. Using FastME 2.1.6.1 (37), TYGS also provides a genome-scale phylogeny based on the Genome BLAST Distance Phylogeny method (GBDP), including the query strains and the automatically selected closest type strains, providing branch support values and a treelikeness indicator (32, 33).

### ORTHOLOGY INFERENCE AND INTERSPECIFIC *Enterococcus* spp. PHYLOGENY

To evaluate the population structure of *E. faecalis* within a broader evolutionary framework, we selected 118 genomes for phylogenetic analysis. This dataset included 55 *E. faecalis* genomes representing the species’ genetic diversity within the scope of this study, four well-characterized *E. faecalis* reference genomes (OG1RF, V583, T5), including the type strain (ATCC 19433 = NBRC 100480), 58 genomes representing other validly published *Enterococcus* species (as of June 13, 2024), and the *Vagococcus fluvialis* DSM 5731 genome as an outgroup. The selection of *E. faecalis* genomes was based on initial genomic and taxonomic analyses, which are detailed in the Results section. Accession numbers, strain names, and taxonomic information for all 59 non-*E. faecalis* genomes are provided in **S2 Dataset**. Of these, 54 were type strains available in GenBank as of the search date, with exceptions also noted in S2 Dataset.

To identify orthologous gene groups (orthogroups), we used OrthoFinder (v2.5.5) with the Multiple Sequence Alignment (MSA) option (“-M msa”) across the 118 genomes (38, 39). OrthoFinder’s relaxed approach, which allows for the inclusion of single-copy orthogroups present in the majority of genomes rather than strictly in all genomes, is particularly suited for highly divergent species (38, 39). This relaxed data criterion improves the phylogenetic signal by incorporating genes that, while not universally present, are conserved in a large proportion of genomes [Further details on the basis for this approach are described in method’s paper (39)]. Consequently, 751 orthogroups were selected, each containing single-copy genes present in at least 95.8% of the genomes. By default, the concatenated MSA of these orthogroups was then generated using MAFFT-linsi (40).

The maximum-likelihood phylogenetic tree was constructed using IQ-TREE v2.3.5 (41). ModelFinder Plus (42) was employed to automatically determine the best-fit amino acid substitution model (LG+F+I+R9), via the “-m MFP” option in IQ-TREE. Branch support was assessed with 1,000 ultrafast bootstrap replicates with UFBoot2 (43). The resulting phylogeny was visualized and customized using iTOL (44).

### *E. faecalis* PANGENOME ANALYSIS AND INTRASPECIFIC PHYLOGENY

Pangenome analysis of the 55 *E. faecalis* genomes, representing the species’ genetic diversity within the scope of this study (see **S1 Dataset**), was conducted using Roary v3.13.0 (45). Out of the 9,950 orthologous gene clusters identified, 1,650 were predicted as core genes (i.e., present in all genomes), and these were then used to build a core gene alignment with MAFFT v7.477 (40). Single nucleotide polymorphisms (SNPs) were extracted from the alignment using SNP-sites v2.5.1 (46), and then used as input for maximum-likelihood tree inference by IQ-TREE (41) under the generalized time reversible (GTR) model of nucleotide substitution with the Gamma model of rate heterogeneity. Branch support was assessed with 10,000 ultrafast bootstrap replicates using UFBoot2 (43). The resulting phylogeny was rooted at the midpoint and customized using iTOL (44).

### ANALYSIS OF CLADE-LEVEL ENRICHED GENES

Based on the *E. faecalis* pangenome analysis described in the previous topic, we used Scoary v1.6.16 (47) to identify genes whose frequencies differed significantly between two groups, hereafter referred to as “subspecies A” and “subspecies B”—arbitrary labels employed in this study solely to facilitate comparisons—indicating enrichment or depletion in either group. Genes were considered significantly associated with a given subspecies if they had a P-value of less than 0.05 after applying the Benjamini-Hochberg correction for multiple comparisons (47). The genes enriched in each subspecies were then classified into their respective COG functional categories (48, 49) using COGClassifier v1.0.5 (50) for a broader functional comparison approach. To further assess differences in the distribution of functional categories between subspecies, we applied the chi-squared test, with statistical significance defined as P-value < 0.01.

### ANALYSIS OF NICHE-SPECIFYING GENES

To identify candidate genes potentially involved in niche differentiation between *E. faecalis* subspecies, we filtered gene clusters that were consistently present in all genomes of one subspecies while absent in the other. These clade-specific genes likely encode functions central to the ecological adaptation of the subspecies, differentiating them from their nearest relatives (17, 51).

Due to the frequent occurrence of inaccurate or incomplete annotations in these genes, initially annotated using Prokka v1.14.5 (25), we performed a secondary functional analysis on representative protein sequences from each clade-specific gene cluster, as identified in the Roary output files. Representative protein sequences were re-annotated through BLASTp (52) searches against the UniProtKB database (53), including reference proteomes and entries from TrEMBL and Swiss-Prot (as of April 2023). Homology was inferred based on sequence similarity, with a cutoff of E-value < 1e-6 (54).

Additionally, protein sequences were further characterized using the InterProScan tool (55), which facilitated the identification of conserved domains, active sites, and other protein signatures of known biological function, thus providing insights into potential functional roles of these niche-specifying genes.

## DATA AVAILABILITY

The raw sequence reads for the local *E. faecalis* strains have been deposited in the NCBI Sequence Read Archive (SRA) under the BioProject accession numbers PRJNA1186978, PRJNA695567, and PRJNA503970. Genome assemblies analyzed in this study include these local strains and publicly available assemblies from GenBank, with accession numbers provided in the S1 and S2 Datasets.

## RESULTS

### DETECTION OF *E. faecalis* STRAINS LACKING CRISPR2: GENETIC DIVERSITY, ISOLATION SOURCES AND CRISPR SYSTEM PROFILES

We detected the presence of CRISPR2 loci in the vast majority of *E. faecalis* strains analyzed in this study (**Fig. 1**). All 71 draft genomes from our collection, representing isolates across 16 distinct STs, were CRISPR2-positive based on both *in silico* PCR screening and CRISPRCasFinder predictions. Similarly, CRISPR2 was identified by at least one of these methodologies in 98.3% (1439 out of 1464) of the *E. faecalis* genomes obtained from GenBank. These publicly available genomes corresponded to 208 distinct STs registered in the PubMLST database (see **S1 Dataset**).

**Fig. 1.**
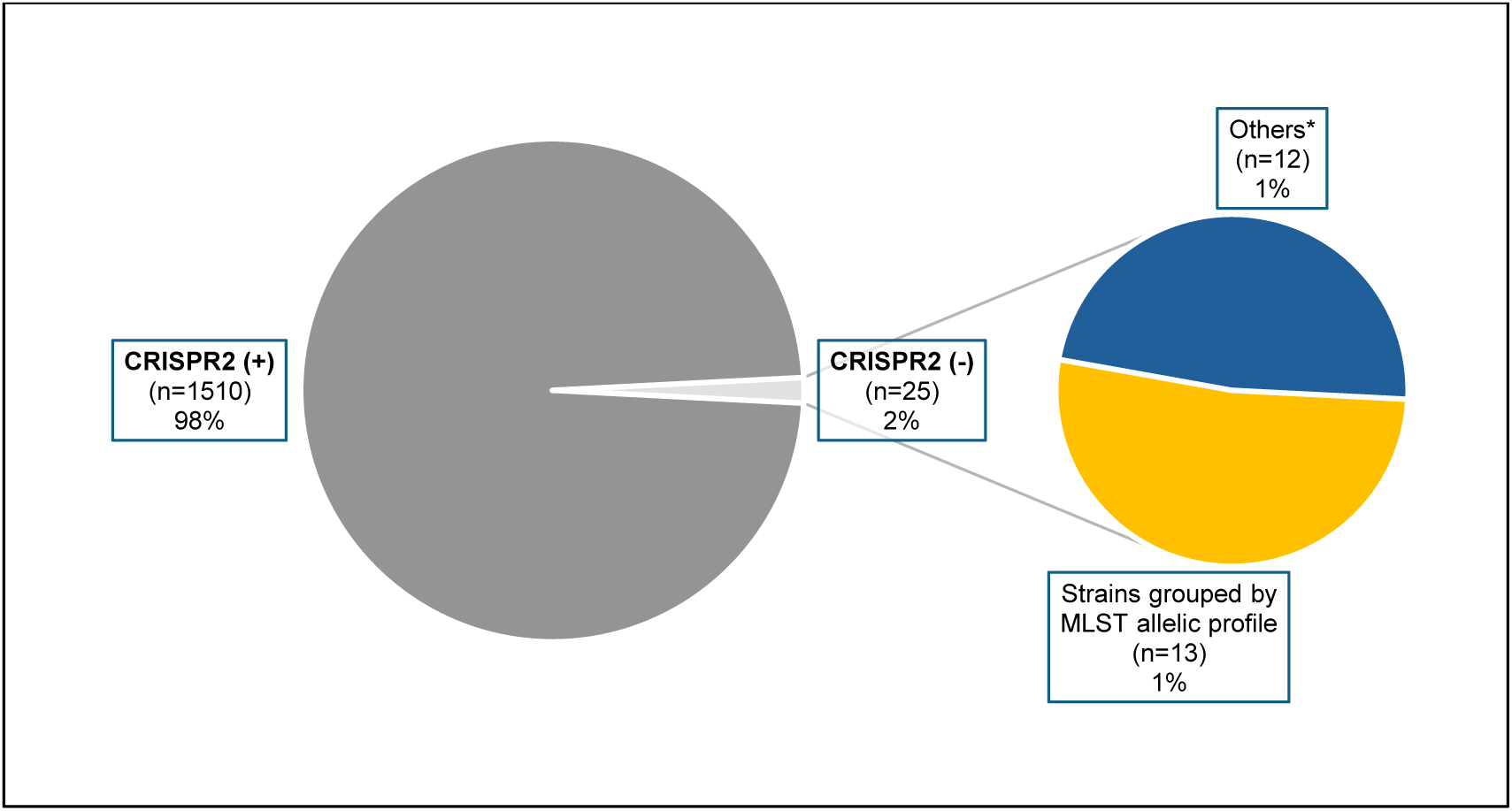
CRISPR2 status in *E. faecalis* genomic samples. The main pie chart categorizes the total genomes (n=1535) into CRISPR2-positive and CRISPR2-negative. The secondary pie chart further classifies CRISPR2-negative genomes into MLST allelic profile-related strains and others. **\***The category “Others” includes genomes with no evident close relationship within the group based on MLST allelic profiles; genomes with statistics indicating low-quality sequencing or assembly; and genomes removed from RefSeq due to various inconsistencies.

Interestingly, out of the remaining 25 genomes where CRISPR2 was not detected by any method, 13 were genetically related, sharing MLST alleles for specific genes. The majority of these were classified as members of ST228, followed by ST624, ST1468, and one strain with an allelic profile not yet registered in the PubMLST database (as of October 2024), which we refer to as “STx” for the purpose of this study (see **Table 1**).

**Table 1.** Sampling details and MLST allelic profiles of the 16 *Enterococcus faecalis* strains lacking CRISPR2.

| Strain | ST | MLST allelic profile |  |  |  |  |  |  | Isolation source | Country | Year of isolation | GenBank assembly accession |
| --- | --- | --- | --- | --- | --- | --- | --- | --- | --- | --- | --- | --- |
|  |  | *gdh* | *gyd* | *pstS* | *gki* | *aroE* | *xpt* | *yqiL* |  |  |  |  |
| CVM N52662 | 228 | 45 | 5 | 44 | 53 | 48 | 41 | 42 | Pork chop | United States | 2014 | GCA_002946755.1 |
| CVM N52467 |  |  |  |  |  |  |  |  | Ground beef | United States | 2014 | GCA_002947085.1 |
| CVM N55265 |  |  |  |  |  |  |  |  | Pork chop | United States | 2014 | GCA_002947435.1 |
| CVM N54548 |  |  |  |  |  |  |  |  | Ground turkey | United States | 2014 | GCA_002948035.1 |
| CVM N53420 |  |  |  |  |  |  |  |  | Pork chop | United States | 2014 | GCA_002948515.1 |
| EN24 |  |  |  |  |  |  |  |  | Chicken carcass rinse samples | New Zealand | 2006 | GCA_004126865.1 |
| C144 |  |  |  |  |  |  |  |  | Beef plant conveyor belts | Canada | 2016 | GCA_006541075.1 |
| C146 |  |  |  |  |  |  |  |  | Beef plant conveyor belts | Canada | 2016 | GCA_006541305.1 |
| C138 |  |  |  |  |  |  |  |  | Beef plant conveyor belts | Canada | 2016 | GCA_006541395.1 |
| R30 |  |  |  |  |  |  |  |  | Retail ground beef | Canada | 2015 | GCA_006541705.1 |
| C116 |  |  |  |  |  |  |  |  | Beef processing facility | Canada | 2015 | GCA_017942395.1 |
| R48 |  |  |  |  |  |  |  |  | Beef processing facility | Canada | 2015 | GCA_017939215.1 |
| 209EA1 | 624 | 59 | 5 | 78 | 53 | 82 | 36 | 42 | Breast meat (*Gallus gallus domesticus*) | United States | unknown | GCA_003319425.1 |
| DSM111623 |  |  |  |  |  |  |  |  | Mouse gastrointestinal tract | Germany | 2019 | GCA_932751055.1 |
| G81 | 1468 | 59 | 50 | 78 | 53 | 82 | 36 | 42 | Beef plant ground product | Canada | 2015 | GCA_006541905.1 |
| CVM N52587 | STx | 45 | 5 | NF | 53 | 48 | 41 | 42 | Ground Beef | United States | 2014 | GCA_002947015.1 |

NF: not found an exact match for the respective locus in the PubMLST database.

The remaining 12 genomes presented several inconsistencies that cast doubt on the reliability of CRISPR2 absence. These genomes were genetically unrelated to each other and to the group of 13 closely related strains, as determined by their distinct MLST allelic profiles (see S1 Dataset). Moreover, many exhibited issues such as low sequencing or assembly quality, inconclusive CRISPR predictions, or had been flagged for removal from RefSeq due to various discrepancies. Given these concerns, it is highly likely that the absence of CRISPR2 in these genomes was an artifact of poor-quality data rather than a true biological feature. Consequently, we excluded these 12 genomes from further analysis and focused exclusively on the 13 genetically related strains that consistently lacked CRISPR2 based on results of both screening methods.

To expand our dataset and investigate these CRISPR2-negative strains further, we conducted a targeted search for additional genomes deposited in GenBank after our initial analysis. This search focused on genomes from the same sequence types (ST228, ST624, ST1468, and STx) or closely related allelic profiles. As a result, three newly deposited genomes belonging to ST228 and ST624 were identified, which also lacked the CRISPR2 locus (**Table 1**). Thus, our final dataset comprised 16 CRISPR2-negative genomes, which became the focus of our study.

Sample details for this genetically related cluster of CRISPR2-negative strains are presented in Table 1. All 16 strains shared the same allelic variants for the housekeeping genes *gki* (53) and *yqiL* (42), with 15 of them (except the ST1468 strain) also sharing the *gyd* (5) allele. Interestingly, most of these strains were isolated from meat products or surfaces in animal processing facilities. Additionally, one strain was isolated from the gastrointestinal tract of a mouse. The strains originated from diverse geographical locations, spanning three continents, and were isolated between 2006 and 2019. Notably, besides lacking the orphan CRISPR2 locus, all 16 strains also lacked the CRISPR1-Cas system but harbored the CRISPR3-Cas system.

### GENOMIC EVIDENCE FOR A DISTINCT *E. faecalis* SUBSPECIES: TAXONOMIC AND PHYLOGENETIC ASSESSMENT

Motivated by the notable absence of CRISPR2 in this genetically related cluster of *E. faecalis* strains, we performed a comprehensive genome-wide evaluation. This analysis sought to clarify their taxonomic classification and determine their position within the broader evolutionary context of the species and its closely related taxa.

For this purpose, we first used the Type (Strain) Genome Server (TYGS) (32), which automatically detects the closest related type strains for each user-submitted genome by default, performing pairwise whole-genome comparisons via the Genome BLAST Distance Phylogeny method (GBDP). GBDP distances are employed to infer phylogenetic trees and to calculate digital DNA:DNA hybridization (dDDH) values, enabling the taxonomic delineation of species and subspecies based on established thresholds (≥70% dDDH for species and ≥79% dDDH for subspecies) (32). As a result, we found that while all 16 CRISPR2-lacking genomes were confirmed as *E. faecalis*, they were classified as a unique subspecies, distinct from the one assigned to their closest type strain (*E. faecalis* ATCC 19433 = NBRC 100480). The TYGS analysis job summary, including taxonomic identification results and pairwise dDDH values between the queried genomes and selected type strains, is provided as supplementary information (**Additional file S1**). A simplified version of this finding is shown in **Fig. 2A**, in which redundant strains were excluded and strain 209EA1 was selected as the representative genome for the CRISPR2-negative cluster.

**Fig. 2.**
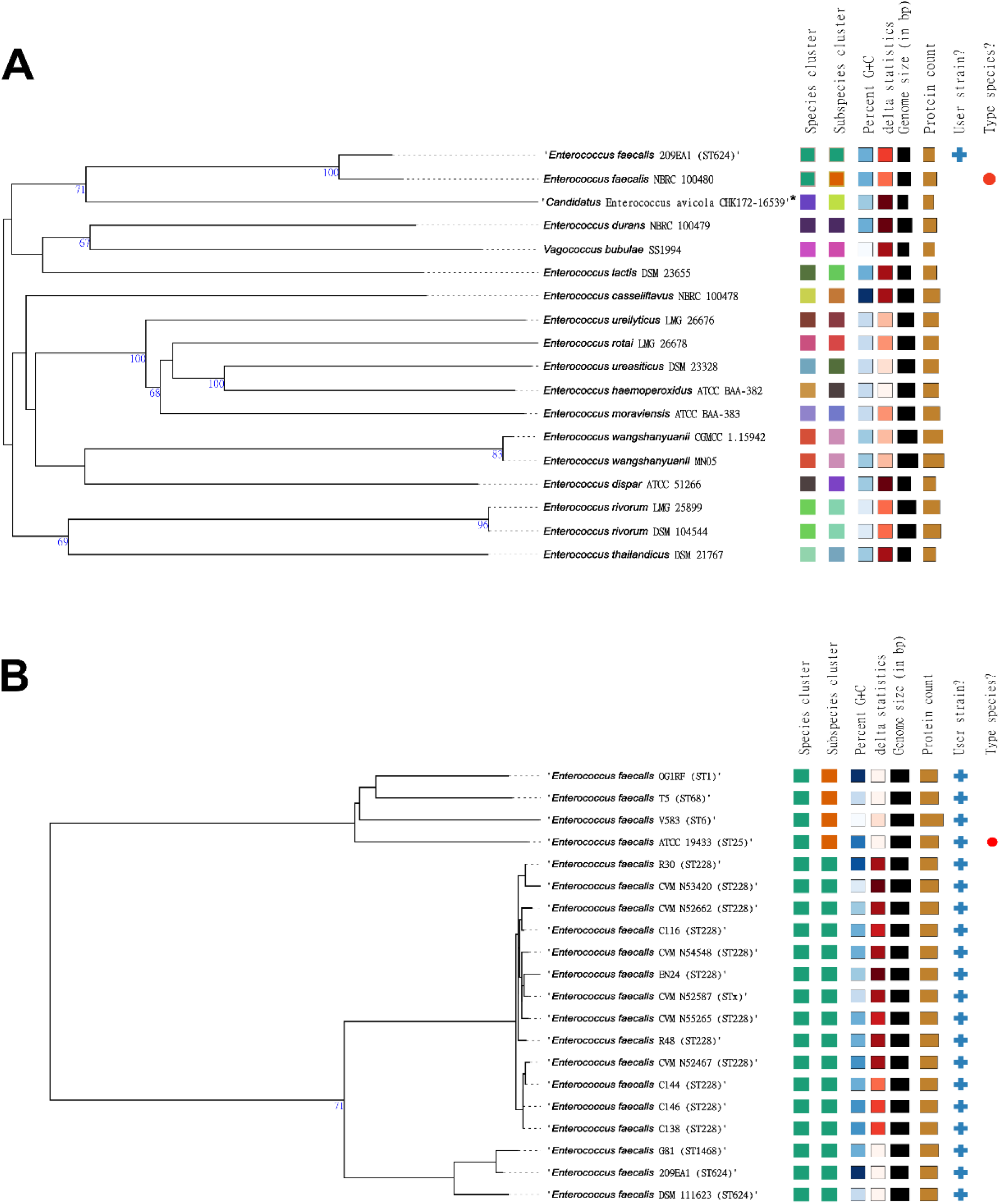
Species and subspecies delineation based on dDDH thresholds. Phylogenetic trees were inferred using FastME 2.1.6.1 from GBDP distances calculated from genome sequences. Branch lengths are scaled according to the GBDP distance formula d5. The numbers above branches represent GBDP pseudo-bootstrap support values > 60% from 100 replications. The trees were midpoint-rooted. The taxonomic classifications at the species and subspecies levels are represented by color codes on the right side of each tree, based on the dDDH values obtained and the established thresholds (≥70% dDDH for species and ≥79% dDDH for subspecies) (**A**) Whole-genome sequence-based phylogeny of *Enterococcus faecalis* 209EA1 (representative strain of the genetically related group of *E. faecalis* lacking CRISPR2) and its closest related type-strains, as determined by TYGS by default. (**B**) Phylogeny focused exclusively on query genomes, encompassing all 16 *E. faecalis* strains lacking CRISPR2 along with *E. faecalis* reference strains, including the type strain ATCC 19433. Sequence types (STs) of each strain are shown in parentheses. **\***Type strain of a proposed species with nomenclatural status not yet validly published.

To gain novel insights into the unprecedented subspecies differentiation of *Enterococcus faecalis*, we conducted a new TYGS analysis. This analysis incorporated additional clonally distinct *E. faecalis* reference strains, including the well-studied model strains OG1RF [derived from the commensal and naturally antibiotic-sensitive human oral isolate OG1 (56)] and V583 [a hospital-adapted VRE isolate (57)], along with the species’ reference genome from NCBI datasets (strain T5). We also included the genome of *E. faecalis*’ type strain. In this analysis, pairwise comparisons were confined to this set of genomes and the 16 CRISPR2-lacking strains. Notably, despite their genetic diversity, all four *E. faecalis* reference strains were grouped within the same subspecies and formed a clade separated from the one containing the 16 CRISPR2-lacking strains (**Fig. 2B**).

For clarity and ease of comparison throughout this study, we will refer to this newly identified subspecies as “subspecies B”, while individuals classified within the most predominant and diverse subspecies (CRISPR2-positive) will be referred to as “subspecies A”.

We expanded our TYGS analysis to investigate whether subspecies B was exclusively composed of genomes lacking CRISPR2, and if the inclusion of additional *E. faecalis* representatives from diverse lineages could reveal new taxonomic subdivisions at the subspecies level. For this purpose, in addition to subspecies B genomes, we incorporated 39 genomes representing 28 distinct STs, encompassing both the genetic diversity from our local collection and other lineages known to harbor CRISPR3-*cas* systems, a trait consistently found in subspecies B (see **S1 Dataset**). The analysis revealed that all 39 genomes were classified within subspecies A (see **Additional file S2**), reinforcing the absence of CRISPR2 as a presumptive feature of subspecies B. Based on these results, we established this set of 39 genomes as the representative subset of subspecies A for subsequent comparative genomic analyses against subspecies B, providing a comprehensive framework for capturing the genetic diversity of *E. faecalis*.

To further validate the reliability of this cladogenesis event and the associated taxonomic separation, we conducted additional phylogenomic reconstructions using different genome sets and approaches. These reconstructions spanned both broad and short evolutionary time scales, encompassing interspecific and intraspecific relationships, respectively (the latter will be presented in the subsequent section). By doing so, we were able to capture different levels of phylogenetic signal, inferred from the concatenated multiple sequence alignments (MSA) of the orthologous groups conserved within each genome set.

Initially, to contextualize our findings within the broader evolutionary landscape of *Enterococcus*, we gathered a dataset of 117 genomes, including 59 *E. faecalis* strains (39 subspecies A, 16 subspecies B, and four additional reference genomes) and 58 unique representatives of other validly published *Enterococcus* species, primarily type strains. In cases where type strain genomes were unavailable or of lower quality than the species’ assigned reference genome in NCBI, we prioritized the highest quality genomes (**S2 Dataset**). *Vagococcus fluvialis* DSM 5731 was selected as the outgroup. Phylogenetic placement was based on the protein sequences of 751 orthologous groups, with a minimum of 95.8% of the genomes (out of the 118 included) containing single-copy genes in any orthologous group (for details, see Materials and Methods section).

The resulting phylogenetic tree corroborated the classification of *E. faecalis* subspecies A and B as a single taxonomic species, forming a well-supported clade distinctly separated by a deep branch from the nearest related *Enterococcus* species (**Fig. 3**). Despite the long-term evolutionary history depicted in the tree, the reconstruction effectively resolved the subspecies-level differentiation within *E. faecalis*, producing two well-supported subclades. Each subclade clusters strains classified in the same subspecies based on dDDH thresholds, demonstrating consistency between recent diversification events and taxonomic separation.

**Fig. 3.**
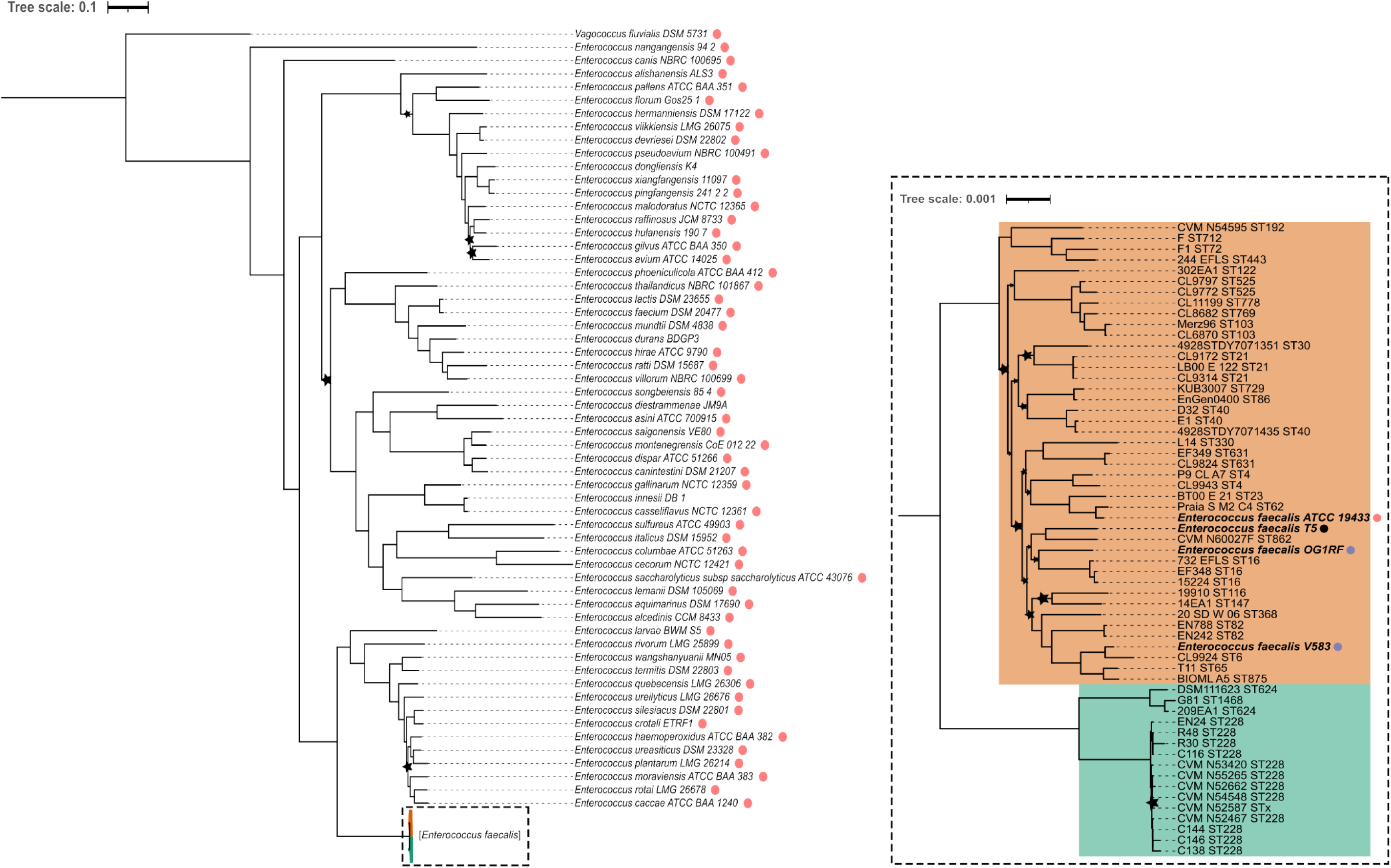
Comprehensive phylogeny of the *Enterococcus* genus illustrating subspecies-level cladogenesis of *E. faecalis*. The tree was estimated using IQ-TREE under the LG+F+I+R9 substitution model with 1,000 bootstrap replicates, based on the concatenation and alignment of 751 protein sequences corresponding to single-copy orthologs present in 95.8% of the 117 sampled enterococci and the outgroup strain *Vagococcus fluvialis* DSM 5731. The tree is presented at different scales. The global phylogeny on the left shows relationships between taxonomic species, with *E. faecalis* strains collapsed into their respective subspecies clades, which are highlighted by colored regions enclosed in a dashed rectangle. On the right, the *E. faecalis* clade is detailed at a fine scale, highlighting subspecies clusters (subspecies A in light orange and subspecies B in light teal). Red dots represent type strains; blue dots indicate *E. faecalis* model strains V583 and OG1RF; the black dot marks the *E. faecalis* reference genome from the NCBI datasets. Stars denote nodes with bootstrap support < 95%.

### PANGENOME AND REVERSE ECOLOGY ANALYSES: GENE CONTENT AND PREDICTED ECOLOGICAL DIFFERENCES BETWEEN SUBSPECIES A AND B

The subset of 55 *E. faecalis* strains (39 subspecies A and 16 subspecies B), included in the previous phylogeny (**Fig. 3**), was selected for a pangenome analysis. This analysis aimed to enhance the phylogenetic resolution of the subspecies-level cladogenesis and to identify genetic determinants potentially involved in their ecological differentiation. The pangenome consisted of 9,950 genes, while the core genome comprised 1,650 single-copy genes. Single nucleotide polymorphisms (SNPs) were extracted from the core gene alignment and used to construct the intraspecific phylogenetic tree presented in **Fig. 4**. Additionally, CRISPR content, predicted antimicrobial resistance (AMR) and virulence-associated genes for each genome are indicated.

**Fig. 4.**
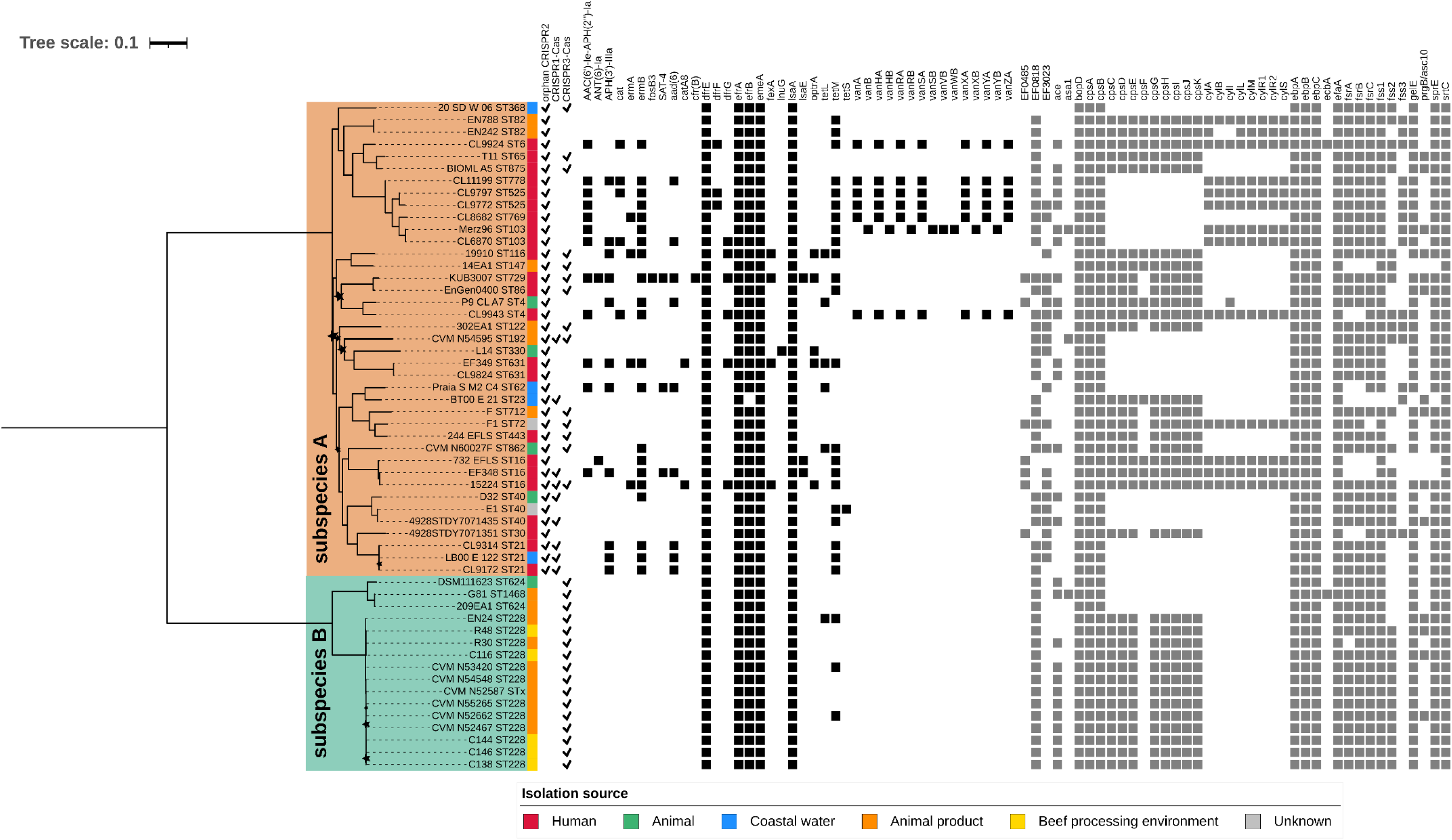
*Enterococcus faecalis* intraspecific phylogeny showcasing the distribution of CRISPR, antimicrobial resistance (AMR), and virulence genes across subspecies A and B. The tree was estimated using IQ-TREE under the GTR+G substitution model with 10,000 bootstrap replicates, based on SNPs extracted from the concatenated alignment of the 1,650 core genes present in all 55 sampled *E. faecalis* genomes. Rooted at the midpoint, the tree depicts *E. faecalis* subspecies A and B as the deepest divergences within the species, shown in colored ranges (subspecies A in light orange and subspecies B in light teal). Stars denote nodes with bootstrap support < 95%. Strain names are followed by their assigned sequence types (STs) in each leaf label. The colored strip adjacent to the strains indicates their respective isolation sources: human (red), animal (green), coastal water (blue), animal product (orange), beef processing environments (yellow), or unknown origin (gray). Screened genetic traits are shown as binary data in the next panels (filled shapes indicate trait presence and omitted shapes indicate absence), with CRISPR loci as checkmarks, antimicrobial resistance genes as black squares, and virulence genes as dark gray squares.

Rooted at the midpoint, the phylogenetic tree reveals a clear subspecies-level divergence within *E. faecalis*, separating into two major clades that reflect the deepest divergences within the species. The larger clade comprising subspecies A genomes displays a more intricate topology with multiple subclades, interconnected by branches of varying lengths and generally lower statistical support. This clade includes most of the STs analyzed, encompassing strains isolated from diverse sources such as humans, animals, animal products, and environmental samples from coastal waters (see **S1 Dataset** for details).

Notably, although CRISPR2 is universally present among genomes of subspecies A, the tree reveals a heterogeneous distribution of complete CRISPR-Cas systems (CRISPR1-Cas and CRISPR3-Cas) within this clade. The subspecies A clade also shows considerable variation in the number of AMR and virulence-associated genes, ranging from genomes with few such genes to strains with prominent multidrug resistance (MDR) profiles, including all vancomycin-resistant enterococci (VRE) isolates.

The clade corresponding to subspecies B was supported by the maximum bootstrap value, corroborating the robustness of this separation. Most strains within this clade, except for ST624 DSM111623, were isolated either directly from animal meat or from surfaces and environments within meat processing facilities, suggesting a common ecological niche. Two well-supported branches are observed, with strains clustering according to their STs. One subclade includes strains from ST228 and its single-locus variant (SLV) STx, with branches exhibiting short lengths, making internal divergence indistinguishable at the current tree scale. The other subclade comprises genomes of the closely related ST624 and ST1468, which are also SLVs of each other.

As previously mentioned, the CRISPR2 and CRISPR1-*cas* loci were consistently absent from the genomes of subspecies B, while CRISPR3-*cas* was universally present across all genomes in this group. In contrast to subspecies A, subspecies B exhibited a more uniform pattern of AMR and virulence genotypes, particularly within the ST228 cluster. The AMR profile of subspecies B was predominantly characterized by genes encoding efflux pumps, such as *drfE, efrA/B,* and *lsaA*, which are associated with MDR and biocide resistance (58, 59). However, these genes were also detected in most subspecies A genomes. Notably, three strains of subspecies B harbored tetracycline resistance genes, specifically *tetM*, with one strain also containing *tetL*.

Although not unique to subspecies B, a broad array of virulence genes was identified in most genomes of this group, especially within the ST228 cluster. These genes include homologues of the *E. faecalis* V583 ORF EF0818, which encodes a family 8 polysaccharide lyase (60); the *bopD* gene, encoding a sugar-binding transcriptional regulator essential for biofilm production (61); the capsule production-associated *cps* operon (except for the *cpsF* gene) (62); the *ebpA/B/C* genes, responsible for encoding endocarditis- and biofilm-associated pili (63); the *efaA* gene, which encodes the *E. faecalis* antigen A (64); the *srtC* gene (also known as *bps*), encoding a biofilm- and pilus-associated sortase (65); the *fss1* gene, encoding a fibrinogen-binding MSCRAMM (Microbial Surface Components Recognizing Adhesive Matrix Molecules) (66); and the *gelE* and *sprE* genes, which encode the secreted proteases gelatinase and serine protease, respectively. These proteases are regulated by the Fsr quorum-sensing system, encoded by the *fsrA/B/C* locus, which was also detected in these genomes (67, 68).

On the other hand, key differences in the resistome and virulome between subspecies A and B were observed. The *ermB* gene and homologues of EF3023 were statistically underrepresented in subspecies B (*P* values < 0.005 after Benjamini-Hochberg correction). The *ermB* gene encodes an enzyme responsible for resistance to macrolides, lincosamides, and streptogramin B (MLS_B_) (69), while EF3023 encodes HylA, a virulence-associated enzyme whose exact role remains to be elucidated (70). Still in the comparative epidemiological context, it is also worth noting that other AMR and virulence genes were exclusively detected in subspecies A. These include determinants involved in high-level aminoglycoside resistance (HLAR) (*aac(6′)-Ie-aph(2′)-Ia*, *ant*(*6*)*-Ia, aph(3’)-IIIa*), glycopeptide resistance (*van* genes), oxazolidinone resistance (*cfr(B)* and *optrA*), as well as the operon encoding the pore-forming exotoxin cytolysin (*cylL_L_*, *cylL_S_*, *cylM*, *cylB*, *cylA*, and *cylI*) and its regulatory genes (*cylR1* and *cylR2*) (71, 72).

Interestingly, the detection of the *ace* gene, which encodes the virulence-associated adhesin to collagen of *E. faecalis* (Ace), revealed subspecies-specific variations in similarity to its corresponding reference sequence in VFDB (NP_814829). Specifically, genes from subspecies A showed higher similarity, as indicated by elevated coverage (100% in most cases) and identity percentages (>95% in all cases) (details are shown in **S3 Dataset**). In contrast, sequences from subspecies B exhibited uniformly lower similarity, with reduced coverage and identity (both indices <85% in all cases) relative to the same reference sequence.

Given the evolutionary distance between subspecies A and B, and considering the generalist lifestyle of *E. faecalis* (6), we aimed to infer the ecological properties that differentiate them as distinct ecotypes based on their genomes. For this purpose, we conducted a pangenome association analysis with the Scoary tool (47), which evaluates components of the pangenome for their associations with specific traits — here, the division between subspecies. This analysis identified 664 genes significantly associated with each subspecies, showing either positive or negative correlations. Of these, 59.3% (394/664) were overrepresented in subspecies A, while the remaining 40.7% (270/664) were more prevalent in subspecies B. The list of subspecies-enriched genes is provided in **S4 Dataset**.

Functional analysis of the genes overrepresented in each subspecies was performed using the COGclassifier tool (50), which classifies prokaryote protein sequences into functional categories based on the COG (Cluster of Orthologous Genes) database. Significant similarity to known sequences was identified for 73.9% (291/394) of the subspecies A-enriched genes and 55.2% (149/270) of the subspecies B-enriched genes (see **S4 Dataset** for details). Although a considerable number of these genes, particularly those associated with subspecies B, could not be categorized into COG functional groups, some notable patterns emerged.

Genes from both subspecies that were successfully classified were distributed across the same 20 COG categories (**Fig. 5A**). Remarkably, subspecies A exhibited a significantly higher proportion of genes likely involved in carbohydrate transport and metabolism (COG category G) compared to subspecies B (*P* = 0.002785) (**Fig. 5B**). The proportions of genes within other COG categories did not show significant variation between the two subspecies.

**Fig. 5.**
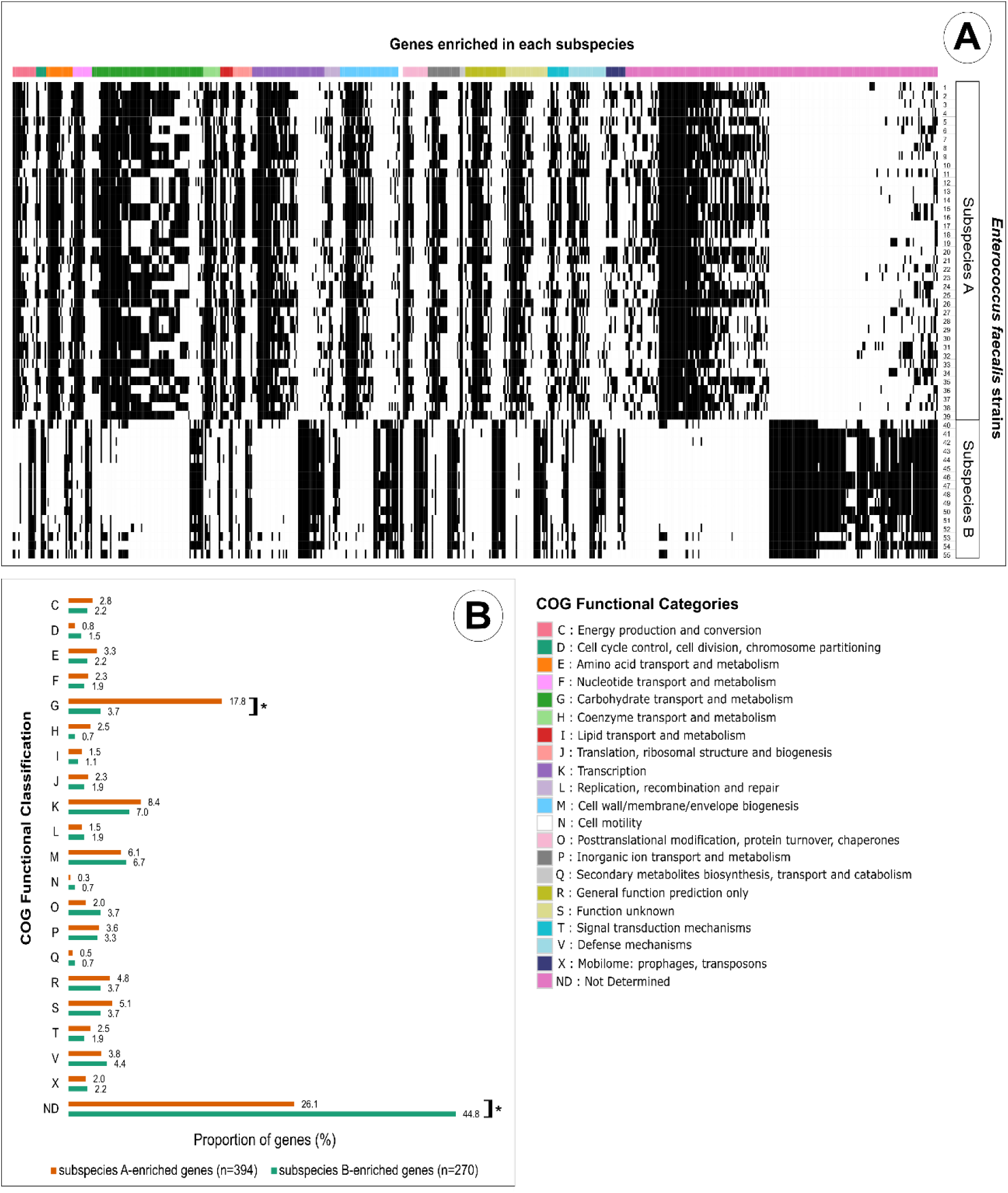
COG functional classification of subspecies-enriched genes. (**A**) Heatmap showing the distribution of the 664 genes significantly associated with subspecies A and B, color-coded by their respective COG functional categories. Black-filled cells represent the presence of a gene in each strain. (**B**) Comparison of gene frequencies across COG functional categories between subspecies A and subspecies B. Each bar represents the proportion of genes annotated in a given functional category in relation to the total number of overrepresented genes in the respective subspecies. An asterisk (*) indicates a P-value <0.01 (chi-squared test).

To identify candidate niche-specifying genes that marked the origin of each subspecies clade (i.e., genes potentially responsible for functional novelties that redefined the ecological niche of their most recent common ancestor) (17, 51, 73), we conducted an additional filtration of the subspecies-enriched genes. Specifically, we focused on those conserved in all members of one clade while absent in all members of the other. This approach led to the preliminary identification of 31 genes associated with subspecies A and 76 genes associated with subspecies B (**S5 Dataset**).

Building on this, we sought to further elucidate the selective pressures that shaped the emergence, maintenance, and diversification of these subspecies by employing a reverse ecology approach (17). This allowed us to infer the likely ecological roles provided by these candidate niche-specifying genes based on their predicted functions. However, the automatic annotation performed by Prokka revealed that 74.2% (23/31) of the candidate niche-specifying genes in subspecies A and 81.6% (62/76) in subspecies B encoded ‘hypothetical proteins,’ indicating a lack of significant matches with known sequences in the databases included in Prokka.

Recognizing these gaps, we conducted a more comprehensive manual annotation of all candidate niche-specifying genes. We utilized BLASTp to compare each sequence against the UniProtKB reference proteomes, Swiss-Prot database, and unreviewed TrEMBL entries, thereby expanding the search coverage. For the follow-up discussion, we focused on the best alignment matches exhibiting significant similarity indicative of homology (see Materials and Methods section for details). Additionally, we examined these sequences against the InterPro protein signature databases to identify protein family memberships, domains, conserved sites, and other features critical for functional characterization.

We observed that 20 genes from subspecies A and 20 from subspecies B exhibited significant similarity to the same reference sequences from *E. faecalis* strain ATCC 700802/V583 in our BLASTp analysis against UniProtKB. Notably, the nuanced differences in sequence similarity between these genes and the reference sequences suggest that they are likely subspecies-specific allelic variants of the same 20 genes, rather than representing 40 distinct orthologous groups (**Table 2**).

**Table 2.**
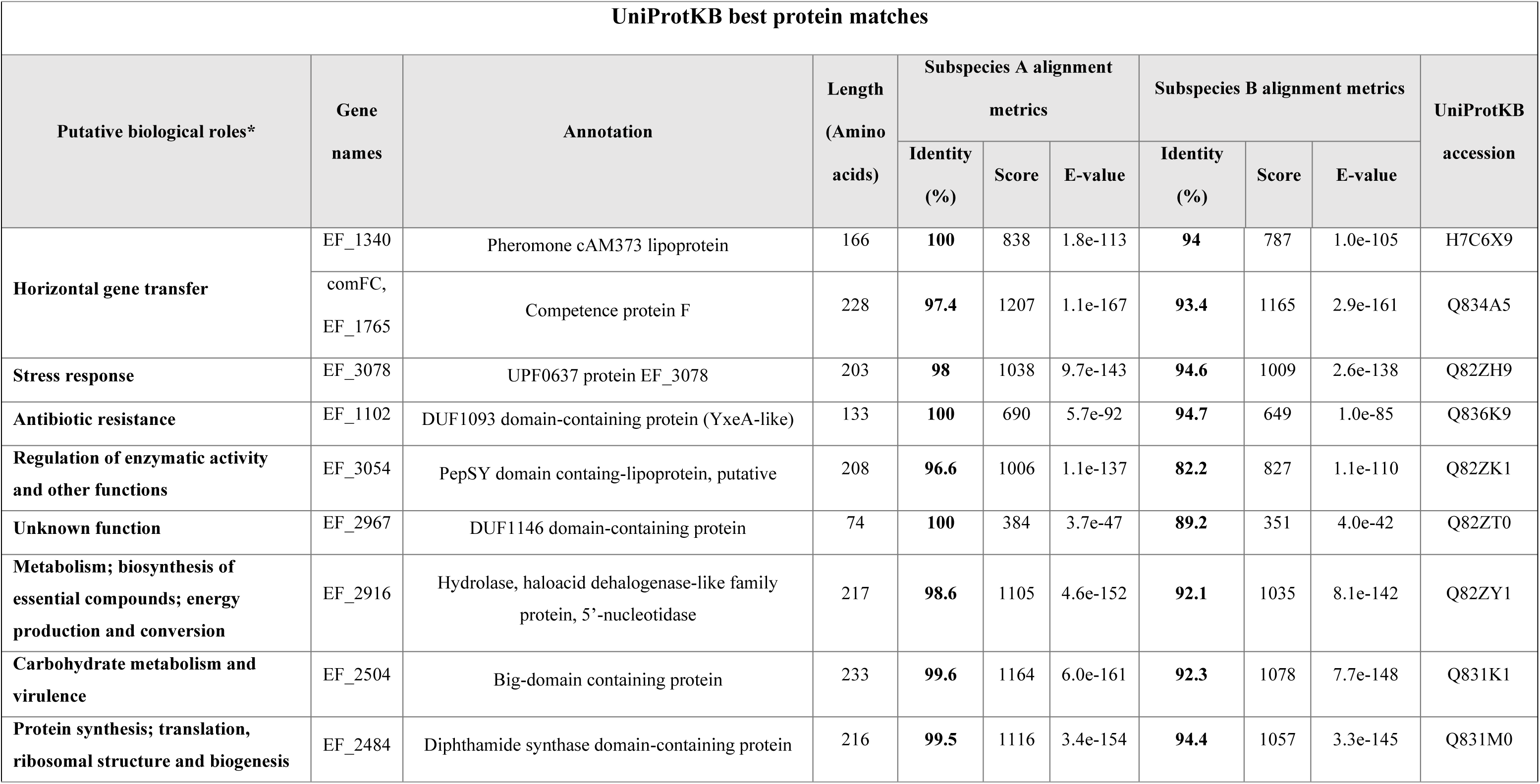
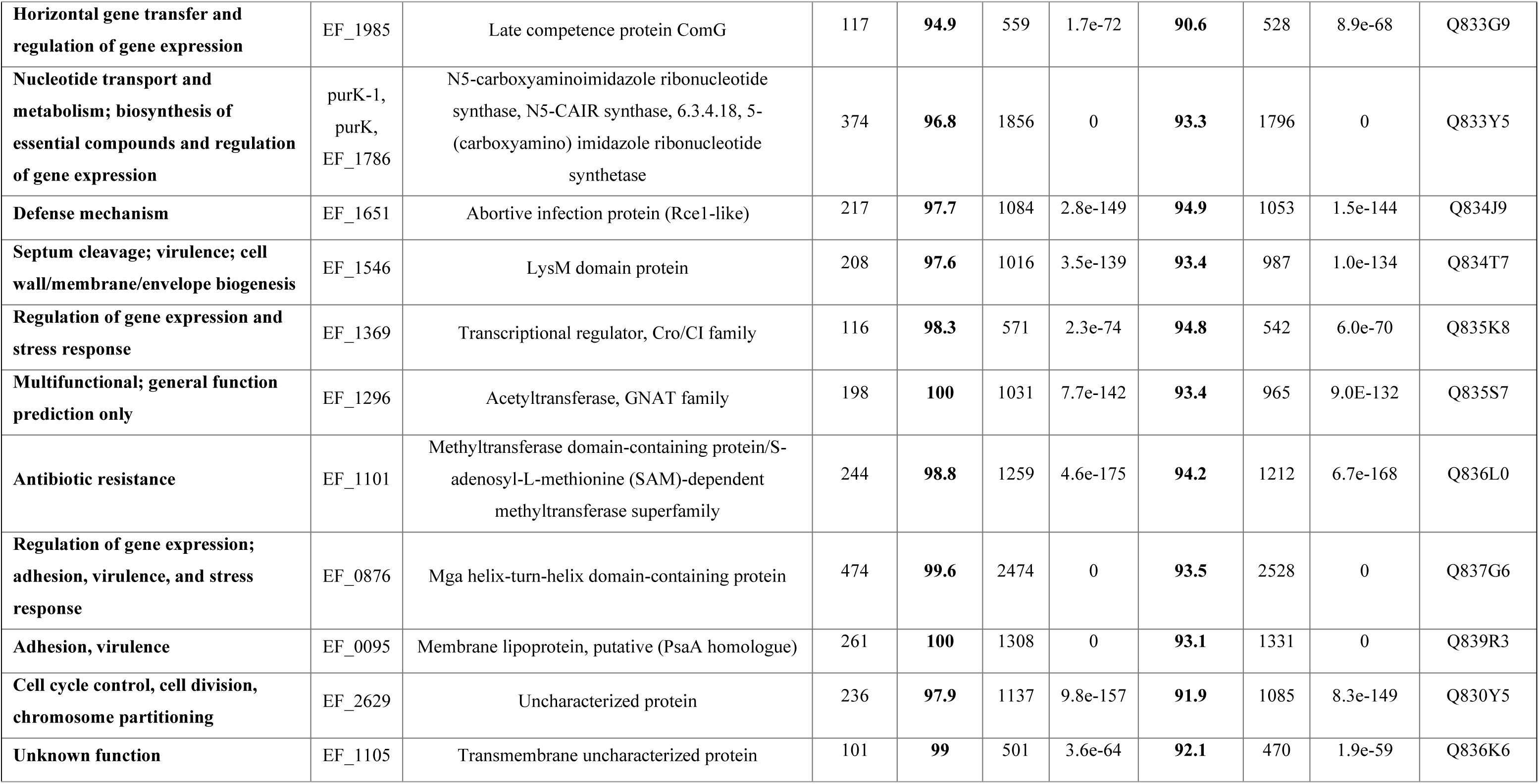
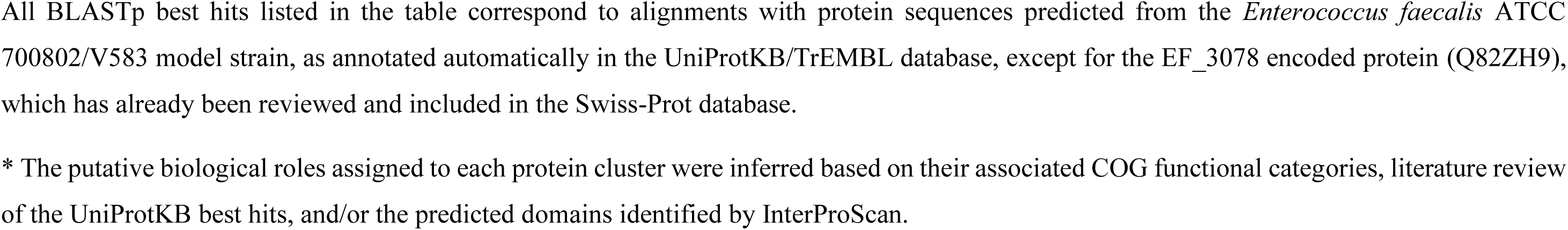
List of *Enterococcus faecalis* subspecies-specific core allelic variants and their putative biological roles based on similarity searches.

| UniProtKB best protein matches |  |  |  |  |  |  |  |  |  |  |
| --- | --- | --- | --- | --- | --- | --- | --- | --- | --- | --- |
| Putative biological roles* | Gene names | Annotation | Length (Amino acids) | Subspecies A alignment metrics |  |  | Subspecies B alignment metrics |  |  | UniProtKB accession |
|  |  |  |  | Identity (%) | Score | E-value | Identity (%) | Score | E-value |  |
| Horizontal gene transfer | EF_1340 | Pheromone cAM373 lipoprotein | 166 | **100** | 838 | 1.8e-113 | **94** | 787 | 1.0e-105 | H7C6X9 |
|  | comFC, EF_1765 | Competence protein F | 228 | **97.4** | 1207 | 1.1e-167 | **93.4** | 1165 | 2.9e-161 | Q834A5 |
| Stress response | EF_3078 | UPF0637 protein EF_3078 | 203 | **98** | 1038 | 9.7e-143 | **94.6** | 1009 | 2.6e-138 | Q82ZH9 |
| Antibiotic resistance | EF_1102 | DUF1093 domain-containing protein (YxeA-like) | 133 | **100** | 690 | 5.7e-92 | **94.7** | 649 | 1.0e-85 | Q836K9 |
| Regulation of enzymatic activity and other functions | EF_3054 | PepSY domain containing-lipoprotein, putative | 208 | **96.6** | 1006 | 1.1e-137 | **82.2** | 827 | 1.1e-110 | Q82ZK1 |
| Unknown function | EF_2967 | DUF1146 domain-containing protein | 74 | **100** | 384 | 3.7e-47 | **89.2** | 351 | 4.0e-42 | Q82ZT0 |
| Metabolism; biosynthesis of essential compounds; energy production and conversion | EF_2916 | Hydrolase, haloacid dehalogenase-like family protein, 5'-nucleotidase | 217 | **98.6** | 1105 | 4.6e-152 | **92.1** | 1035 | 8.1e-142 | Q82ZY1 |
| Carbohydrate metabolism and virulence | EF_2504 | Big-domain containing protein | 233 | **99.6** | 1164 | 6.0e-161 | **92.3** | 1078 | 7.7e-148 | Q831K1 |
| Protein synthesis; translation, ribosomal structure and biogenesis | EF_2484 | Diphthamide synthase domain-containing protein | 216 | **99.5** | 1116 | 3.4e-154 | **94.4** | 1057 | 3.3e-145 | Q831M0 |
| **Horizontal gene transfer and regulation of gene expression** | EF_1985 | Late competence protein ComG | 117 | **94.9** | 559 | 1.7e-72 | **90.6** | 528 | 8.9e-68 | Q833G9 |
| **Nucleotide transport and metabolism; biosynthesis of essential compounds and regulation of gene expression** | purK-1, purK, EF_1786 | N5-carboxyaminoimidazole ribonucleotide synthase, N5-CAIR synthase, 6.3.4.18, 5-(carboxyamino) imidazole ribonucleotide synthetase | 374 | **96.8** | 1856 | 0 | **93.3** | 1796 | 0 | Q833Y5 |
| **Defense mechanism** | EF_1651 | Abortive infection protein (Rce1-like) | 217 | **97.7** | 1084 | 2.8e-149 | **94.9** | 1053 | 1.5e-144 | Q834J9 |
| **Septum cleavage; virulence; cell wall/membrane/envelope biogenesis** | EF_1546 | LysM domain protein | 208 | **97.6** | 1016 | 3.5e-139 | **93.4** | 987 | 1.0e-134 | Q834T7 |
| **Regulation of gene expression and stress response** | EF_1369 | Transcriptional regulator, Cro/CI family | 116 | **98.3** | 571 | 2.3e-74 | **94.8** | 542 | 6.0e-70 | Q835K8 |
| **Multifunctional; general function prediction only** | EF_1296 | Acetyltransferase, GNAT family | 198 | **100** | 1031 | 7.7e-142 | **93.4** | 965 | 9.0E-132 | Q835S7 |
| **Antibiotic resistance** | EF_1101 | Methyltransferase domain-containing protein/S-adenosyl-L-methionine (SAM)-dependent methyltransferase superfamily | 244 | **98.8** | 1259 | 4.6e-175 | **94.2** | 1212 | 6.7e-168 | Q836L0 |
| **Regulation of gene expression; adhesion, virulence, and stress response** | EF_0876 | Mga helix-turn-helix domain-containing protein | 474 | **99.6** | 2474 | 0 | **93.5** | 2528 | 0 | Q837G6 |
| **Adhesion, virulence** | EF_0095 | Membrane lipoprotein, putative (PsaA homologue) | 261 | **100** | 1308 | 0 | **93.1** | 1331 | 0 | Q839R3 |
| **Cell cycle control, cell division, chromosome partitioning** | EF_2629 | Uncharacterized protein | 236 | **97.9** | 1137 | 9.8e-157 | **91.9** | 1085 | 8.3e-149 | Q830Y5 |
| **Unknown function** | EF_1105 | Transmembrane uncharacterized protein | 101 | **99** | 501 | 3.6e-64 | **92.1** | 470 | 1.9e-59 | Q836K6 |

All BLASTp best hits listed in the table correspond to alignments with protein sequences predicted from the *Enterococcus faecalis* ATCC 700802/V583 model strain, as annotated automatically in the UniProtKB/TrEMBL database, except for the EF\_3078 encoded protein (Q82ZH9), which has already been reviewed and included in the Swiss-Prot database.
\* The putative biological roles assigned to each protein cluster were inferred based on their associated COG functional categories, literature review of the UniProtKB best hits, and/or the predicted domains identified by InterProScan.

The putative biological functions of the proteins encoded by these subspecies-specific allelic variants are summarized in **Table 2**. These predictions highlight several proteins involved in fundamental cellular processes, including enzymes critical for biosynthesis (e.g., those required for protein synthesis), stress response mechanisms, regulation of gene expression, and energy production. Additionally, some proteins are potentially implicated in horizontal gene transfer, defense mechanisms, virulence, or possess currently unknown functions.

Furthermore, additional 21 candidate niche-specifying genes from subspecies B also exhibited significant similarity to sequences from *E. faecalis* ATCC 700802/V583, a member of subspecies A, as illustrated in **Fig. 2B** (see **S5 Dataset** for further details on these genes and the best alignment matches for their encoded proteins). To mitigate potential sampling biases, we focused our analysis on candidate niche-specifying genes that did not exhibit significant similarity to sequences from the opposing subspecies (see **S5 Dataset**). This approach ensures that the genes analyzed are more likely to be truly subspecies-specific orthologs.

Following this stringent filtering process, we identified 11 orthologous groups unique to subspecies A strains and 35 unique to subspecies B strains. Detailed information on the proteins encoded by these niche-specifying genes is provided in **Tables 3** and **4**, respectively. The implications of these findings are discussed in the following section.

**Table 3.** List of *Enterococcus faecalis* subspecies A-specific proteins and their predicted biological roles based on similarity searches.

| UniProtKB best protein matches |  |  |  |  |  |  |  |
| --- | --- | --- | --- | --- | --- | --- | --- |
| Putative biological roles* | Gene names | Annotation | Length (Amino acids) | Subspecies A alignment metrics |  |  | UniProtKB accession |
|  |  |  |  | Identity (%) | Score | E-value |  |
| Regulation of gene expression | EF_2063 | SoxS-like Transcriptional regulator, AraC family | 287 | **97.6** | 1467 | 0 | Q833A4 |
| Secondary metabolites biosynthesis, transport and catabolism; phosphorus metabolism | EF_1926 | 5-bromo-4-chloroindolyl phosphate hydrolysis protein | 179 | **97.8** | 887 | 1.6e-120 | Q833L9 |
| Unknown function | EF_1925 | Transmembrane uncharacterized protein | 281 | **99.3** | 1457 | 0 | Q833M0 |
| Transport and binding and energy metabolism | dcuC, EF_1920 | C4-dicarboxylate anaerobic carrier DcuC | 457 | **99.8** | 2221 | 0 | Q833M4 |
| Amino acid transport and metabolism; stress response and antibiotic resistance | EF_1919 | Acetyltransferase, GNAT family | 162 | **100** | 851 | 1.4e-115 | Q833M5 |
| Cell wall/membrane/envelope biogenesis; transport and antibiotic resistance | EF_1672 | Peptide ABC superfamily ATP binding cassette transporter permease | 427 | **99.8** | 2076 | 0 | Q834I0 |
| Carbohydrate transport and metabolism; virulence | scrB-1, EF_1603 | Sucrose-6-phosphate hydrolase, 3.2.1.26, Invertase | 486 | **98.8** | 2566 | 0 | Q834P0 |
| Carbohydrate transport and metabolism; virulence | EF_1602 | Glycosyl hydrolase, family 13 (associated with sucrose uptake PTS-system) | 557 | **99.6** | 2970 | 0 | Q834P1 |
| **Secondary metabolites biosynthesis, transport and catabolism; stress response** | EF_1138 | Oxidoreductase, aldo/keto reductase family | 274 | **99.3** | 1427 | 0 | Q836H3 |
| **Unknown function** | EF_1137 | Membrane uncharacterized protein | 91 | **98.9** | 470 | 3.1e-59 | Q836H4 |
| **Unknown function** | EF_1098 | DUF898 family transmembrane protein | 101 | **100** | 559 | 5.1e-73 | Q836L3 |

All BLASTp best hits listed in the table correspond to alignments with protein sequences predicted from the *Enterococcus faecalis* ATCC 700802/V583 model strain, as annotated automatically in the UniProtKB/TrEMBL database.
\* The putative biological roles assigned to each protein cluster were inferred based on their associated COG functional categories, literature review of the UniProtKB best hits, and/or the predicted domains identified by InterProScan.

**Table 4.**
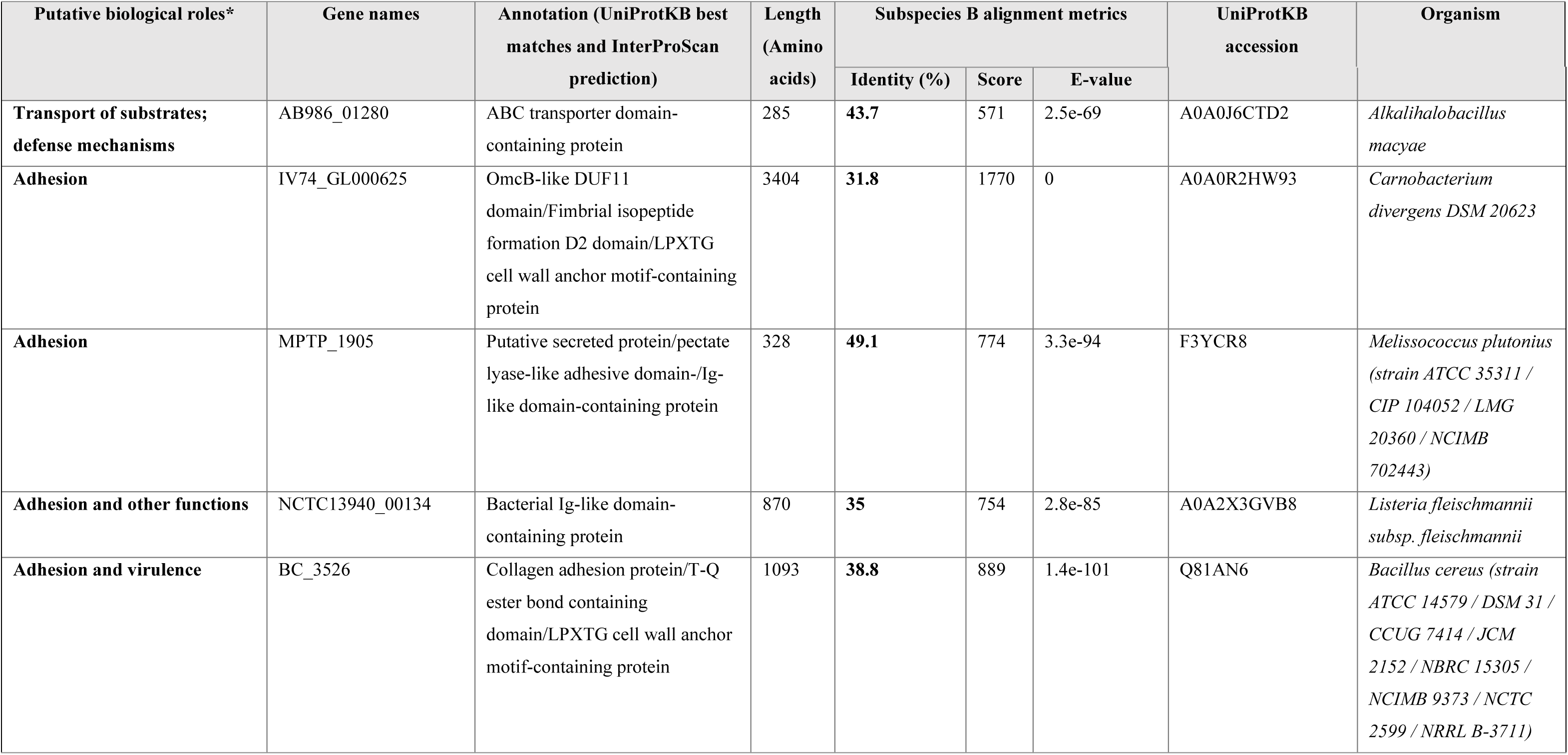
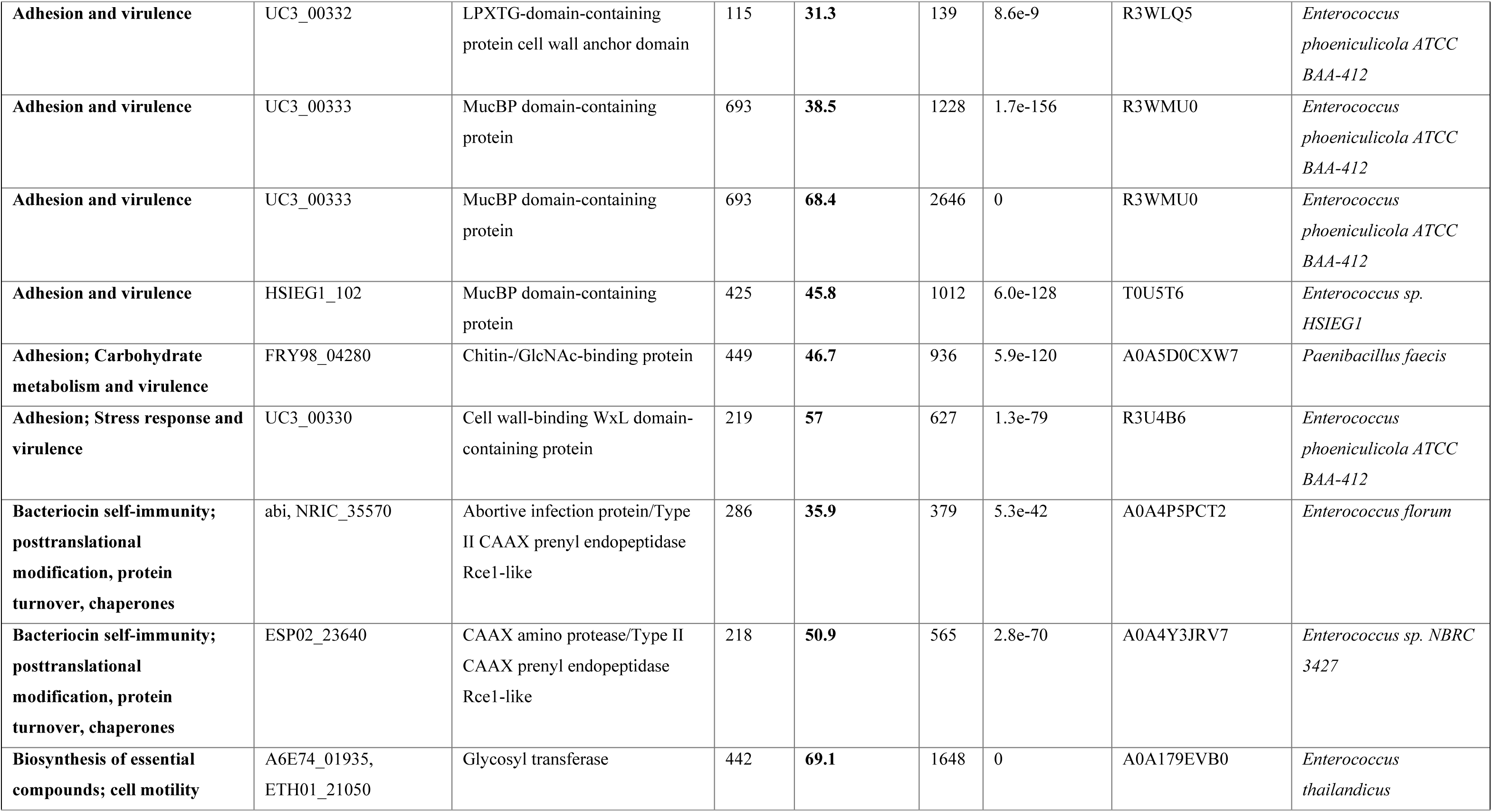
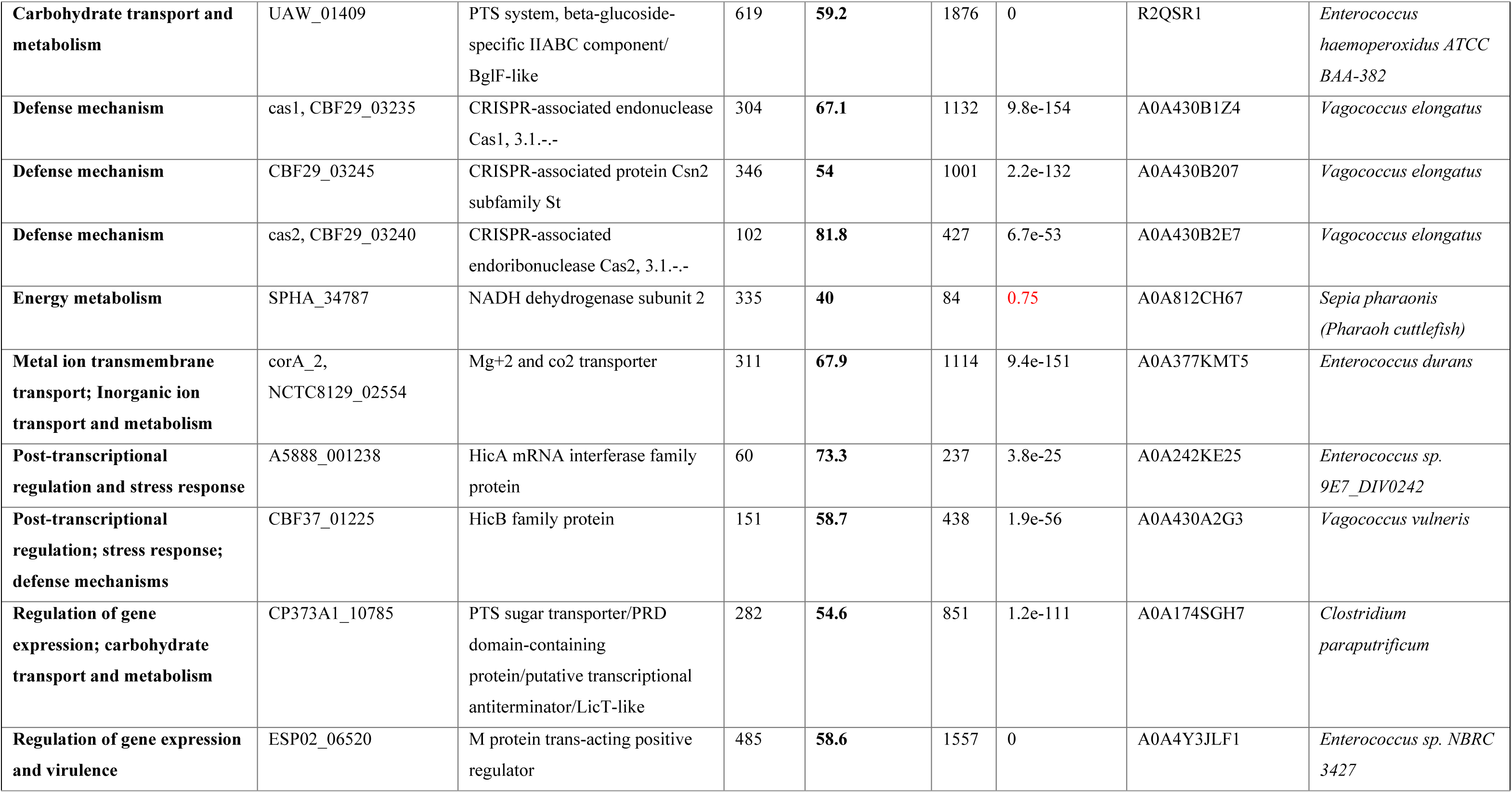
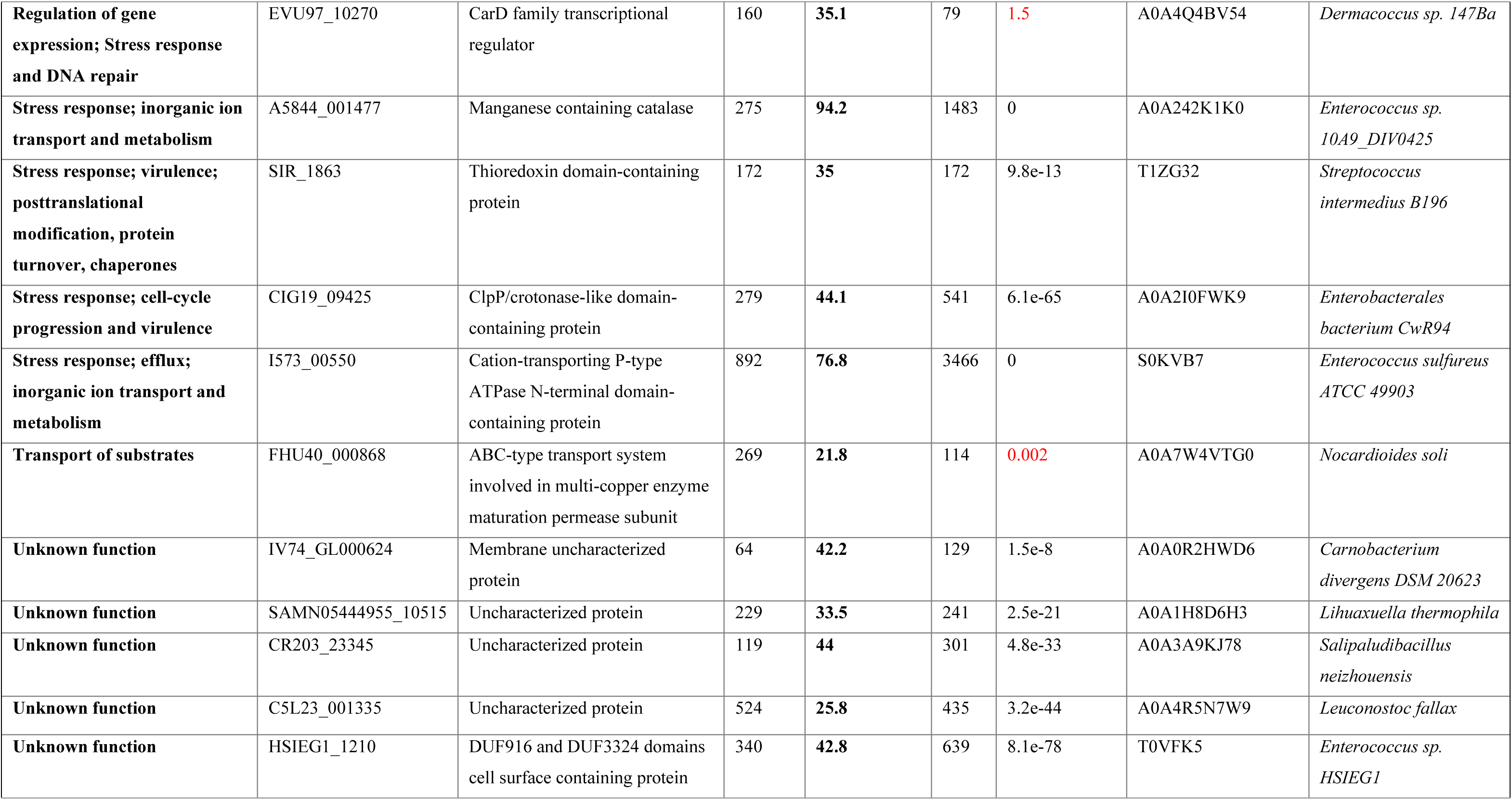
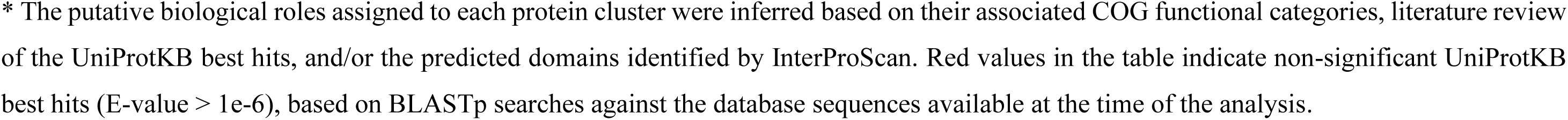
List of *Enterococcus faecalis* subspecies B-specific proteins and their predicted biological roles based on similarity searches.

| Putative biological roles* | Gene names | Annotation (UniProtKB best matches and InterProScan prediction) | Length (Amino acids) | Subspecies B alignment metrics |  |  | UniProtKB accession | Organism |
| --- | --- | --- | --- | --- | --- | --- | --- | --- |
|  |  |  |  | Identity (%) | Score | E-value |  |  |
| Transport of substrates; defense mechanisms | AB986_01280 | ABC transporter domain-containing protein | 285 | 43.7 | 571 | 2.5e-69 | A0A0J6CTD2 | *Alkalihalobacillus macyae* |
| Adhesion | IV74_GL000625 | OmcB-like DUF11 domain/Fimbrial isopeptide formation D2 domain/LPXTG cell wall anchor motif-containing protein | 3404 | 31.8 | 1770 | 0 | A0A0R2HW93 | *Carnobacterium divergens* DSM 20623 |
| Adhesion | MPTP_1905 | Putative secreted protein/pectate lyase-like adhesive domain-/Ig-like domain-containing protein | 328 | 49.1 | 774 | 3.3e-94 | F3YCR8 | *Melissococcus plutonius* (strain ATCC 35311 / CIP 104052 / LMG 20360 / NCIMB 702443) |
| Adhesion and other functions | NCTC13940_00134 | Bacterial Ig-like domain-containing protein | 870 | 35 | 754 | 2.8e-85 | A0A2X3GVB8 | *Listeria fleischmannii* subsp. *fleischmannii* |
| Adhesion and virulence | BC_3526 | Collagen adhesion protein/T-Q ester bond containing domain/LPXTG cell wall anchor motif-containing protein | 1093 | 38.8 | 889 | 1.4e-101 | Q81AN6 | *Bacillus cereus* (strain ATCC 14579 / DSM 31 / CCUG 7414 / JCM 2152 / NBRC 15305 / NCIMB 9373 / NCTC 2599 / NRRL B-3711) |
| **Adhesion and virulence** | UC3_00332 | LPXTG-domain-containing protein cell wall anchor domain | 115 | **31.3** | 139 | 8.6e-9 | R3WLQ5 | *Enterococcus phoeniculicola* ATCC BAA-412 |
| **Adhesion and virulence** | UC3_00333 | MucBP domain-containing protein | 693 | **38.5** | 1228 | 1.7e-156 | R3WMU0 | *Enterococcus phoeniculicola* ATCC BAA-412 |
| **Adhesion and virulence** | UC3_00333 | MucBP domain-containing protein | 693 | **68.4** | 2646 | 0 | R3WMU0 | *Enterococcus phoeniculicola* ATCC BAA-412 |
| **Adhesion and virulence** | HSIEG1_102 | MucBP domain-containing protein | 425 | **45.8** | 1012 | 6.0e-128 | T0U5T6 | *Enterococcus* sp. *HSIEG1* |
| **Adhesion; Carbohydrate metabolism and virulence** | FRY98_04280 | Chitin-/GlcNAc-binding protein | 449 | **46.7** | 936 | 5.9e-120 | A0A5D0CXW7 | *Paenibacillus faecis* |
| **Adhesion; Stress response and virulence** | UC3_00330 | Cell wall-binding WxL domain-containing protein | 219 | **57** | 627 | 1.3e-79 | R3U4B6 | *Enterococcus phoeniculicola* ATCC BAA-412 |
| **Bacteriocin self-immunity; posttranslational modification, protein turnover, chaperones** | abi, NRIC_35570 | Abortive infection protein/Type II CAAX prenyl endopeptidase Rce1-like | 286 | **35.9** | 379 | 5.3e-42 | A0A4P5PCT2 | *Enterococcus florum* |
| **Bacteriocin self-immunity; posttranslational modification, protein turnover, chaperones** | ESP02_23640 | CAAX amino protease/Type II CAAX prenyl endopeptidase Rce1-like | 218 | **50.9** | 565 | 2.8e-70 | A0A4Y3JRV7 | *Enterococcus* sp. *NBRC 3427* |
| **Biosynthesis of essential compounds; cell motility** | A6E74_01935, ETH01_21050 | Glycosyl transferase | 442 | **69.1** | 1648 | 0 | A0A179EVB0 | *Enterococcus thailandicus* |
| **Carbohydrate transport and metabolism** | UAW_01409 | PTS system, beta-glucoside-specific IIABC component/BglF-like | 619 | **59.2** | 1876 | 0 | R2QSR1 | *Enterococcus haemoperoxidus* ATCC BAA-382 |
| **Defense mechanism** | cas1, CBF29_03235 | CRISPR-associated endonuclease Cas1, 3.1.-.- | 304 | **67.1** | 1132 | 9.8e-154 | A0A430B1Z4 | *Vagococcus elongatus* |
| **Defense mechanism** | CBF29_03245 | CRISPR-associated protein Csn2 subfamily St | 346 | **54** | 1001 | 2.2e-132 | A0A430B207 | *Vagococcus elongatus* |
| **Defense mechanism** | cas2, CBF29_03240 | CRISPR-associated endoribonuclease Cas2, 3.1.-.- | 102 | **81.8** | 427 | 6.7e-53 | A0A430B2E7 | *Vagococcus elongatus* |
| **Energy metabolism** | SPHA_34787 | NADH dehydrogenase subunit 2 | 335 | **40** | 84 | **0.75** | A0A812CH67 | *Sepia pharaonis* (Pharaoh cuttlefish) |
| **Metal ion transmembrane transport; Inorganic ion transport and metabolism** | corA_2, NCTC8129_02554 | Mg+2 and co2 transporter | 311 | **67.9** | 1114 | 9.4e-151 | A0A377KMT5 | *Enterococcus durans* |
| **Post-transcriptional regulation and stress response** | A5888_001238 | HicA mRNA interferase family protein | 60 | **73.3** | 237 | 3.8e-25 | A0A242KE25 | *Enterococcus sp.* 9E7_DIV0242 |
| **Post-transcriptional regulation; stress response; defense mechanisms** | CBF37_01225 | HicB family protein | 151 | **58.7** | 438 | 1.9e-56 | A0A430A2G3 | *Vagococcus vulneris* |
| **Regulation of gene expression; carbohydrate transport and metabolism** | CP373A1_10785 | PTS sugar transporter/PRD domain-containing protein/putative transcriptional antiterminator/LicT-like | 282 | **54.6** | 851 | 1.2e-111 | A0A174SGH7 | *Clostridium paraputrificum* |
| **Regulation of gene expression and virulence** | ESP02_06520 | M protein trans-acting positive regulator | 485 | **58.6** | 1557 | 0 | A0A4Y3JLF1 | *Enterococcus sp.* NBRC 3427 |
| **Regulation of gene expression; Stress response and DNA repair** | EVU97_10270 | CarD family transcriptional regulator | 160 | **35.1** | 79 | 1.5 | A0A4Q4BV54 | *Dermacoccus sp. 147Ba* |
| **Stress response; inorganic ion transport and metabolism** | A5844_001477 | Manganese containing catalase | 275 | **94.2** | 1483 | 0 | A0A242K1K0 | *Enterococcus sp. 10A9_DIV0425* |
| **Stress response; virulence; posttranslational modification, protein turnover, chaperones** | SIR_1863 | Thioredoxin domain-containing protein | 172 | **35** | 172 | 9.8e-13 | T1ZG32 | *Streptococcus intermedius B196* |
| **Stress response; cell-cycle progression and virulence** | CIG19_09425 | ClpP/crotonase-like domain-containing protein | 279 | **44.1** | 541 | 6.1e-65 | A0A2I0FWK9 | *Enterobacteriales bacterium CwR94* |
| **Stress response; efflux; inorganic ion transport and metabolism** | I573_00550 | Cation-transporting P-type ATPase N-terminal domain-containing protein | 892 | **76.8** | 3466 | 0 | S0KVB7 | *Enterococcus sulfureus ATCC 49903* |
| **Transport of substrates** | FHU40_000868 | ABC-type transport system involved in multi-copper enzyme maturation permease subunit | 269 | **21.8** | 114 | 0.002 | A0A7W4VTG0 | *Nocardioides soli* |
| **Unknown function** | IV74_GL000624 | Membrane uncharacterized protein | 64 | **42.2** | 129 | 1.5e-8 | A0A0R2HWD6 | *Carnobacterium divergens DSM 20623* |
| **Unknown function** | SAMN05444955_10515 | Uncharacterized protein | 229 | **33.5** | 241 | 2.5e-21 | A0A1H8D6H3 | *Lihuaxuella thermophila* |
| **Unknown function** | CR203_23345 | Uncharacterized protein | 119 | **44** | 301 | 4.8e-33 | A0A3A9KJ78 | *Salipaludibacillus neizhouensis* |
| **Unknown function** | C5L23_001335 | Uncharacterized protein | 524 | **25.8** | 435 | 3.2e-44 | A0A4R5N7W9 | *Leuconostoc fallax* |
| **Unknown function** | HSIEG1_1210 | DUF916 and DUF3324 domains cell surface containing protein | 340 | **42.8** | 639 | 8.1e-78 | T0VFK5 | *Enterococcus sp. HSIEG1* |

\* The putative biological roles assigned to each protein cluster were inferred based on their associated COG functional categories, literature review of the UniProtKB best hits, and/or the predicted domains identified by InterProScan. Red values in the table indicate non-significant UniProtKB best hits (E-value > 1e-6), based on BLASTp searches against the database sequences available at the time of the analysis.

## DISCUSSION

The distinctive biological mechanism of CRISPR systems, particularly their intrinsic ability to sequentially incorporate fragments of foreign DNA, thereby generating a hypervariable region within the host chromosome, places these loci as true biological records. This property renders them valuable for reconstructing historical events, such as host-virus interactions in natural environments, as well as for assessing genetic diversity (74, 75). The potential of CRISPR loci has already been explored for various applications, including tracking clinically relevant lineages and elucidating their origins and evolutionary paths (76, 77). In *E. faecalis*, the orphan CRISPR2 locus has been explored as an alternative target for genetic diversity analysis and phylogenetic studies, enabling clonality predictions and the identification of recombination events among STs (12–14).

Our original goal was to evaluate CRISPR2 array variability for genotyping and phylogenetic analysis of *E. faecalis* strains from diverse sources. However, initial analysis revealed an unexpected finding: a genetically related group of strains lacking CRISPR2. This observation challenges the prevailing assumption that CRISPR2 is ubiquitous, a component of the species’ core genome (8, 12). Given this intriguing observation and the fact that the orphan CRISPR2 locus, in the absence of associated *cas* genes, has no clearly defined biological role, we redirected our focus to explore the implications of CRISPR2 absence in the context of the species’ evolutionary trajectory. Due to the fact that none of the CRISPR2-negative strains were part of our local bacterial culture collection, our investigation was limited to genomic approaches.

Phylogenomic analyses revealed that the CRISPR2-negative strains form one of two *E. faecalis* major clades. Their most recent common ancestor (MRCA) diverged early in the evolutionary history of the species, leading to the independent evolution of these two lineages. These lineages can be classified into distinct subspecies based on dDDH thresholds for taxonomic delineation (referred to here as subspecies A and B, consisting exclusively of CRISPR2-positive and CRISPR2-negative genomes, respectively) (36). Objectively, this means that pairwise genomic comparisons between any representatives of each subspecies reveal hybridization values below 79%. Because no validly published *E. faecalis* child taxa were listed in the LPSN database at the time of writing, we believe this is the first study to describe a subspecies-level division for *E. faecalis*, based on the current criterion used for taxonomic classification of bacteria (33, 34, 78).

Our findings challenge the previous understanding of *E. faecalis* population structure and its generalist nature. By revealing well-supported and distantly related subspecies clades within *E. faecalis*, our results question the assumption that this species lacks a multiclade structure and is devoid of clearly divergent groups, as suggested by earlier studies (6, 15). Given that these prior studies did not include representative STs from subspecies B, it is plausible that their conclusions were influenced by a limited genomic sampling that lacked sufficient genetic diversity. In contrast, our study included a more comprehensive set of lineages, providing a broader perspective on the species’ population structure.

The internal topology of the subspecies A clade reflects its extensive genetic and ecological diversity. This clade includes STs predominantly found in hospital environments, with multidrug resistance profiles and a strong association with HAIs— such as STs 6, 103, 525, and 778, including all VRE isolates in our study (e.g., the well-characterized clinical isolate V583). Additionally, the clade encompasses generalist STs like STs 4, 16, 21, and 40, which are widespread across various ecological niches and display variable CRISPR-Cas content, alongside strains such as the commensal-like OG1RF (ST1), the species type strain ATCC 19433 (ST25), and the NCBI reference genome T5 (ST68) (14, 15, 79–81). The low bootstrap support observed among these lineages suggests complex evolutionary dynamics, likely driven by diverse selective pressures and rapid diversification, which complicate the reconstruction of a clear evolutionary pathway. While this finding is consistent with the aforementioned studies focused on subspecies A, it does not fully capture the population structure of *E. faecalis*, particularly regarding the phylogenetic positioning and evolutionary significance of subspecies B.

On the other hand, the inclusion of subspecies B in the phylogenetic reconstructions of *E. faecalis* revealed significantly divergent lineages, as evidenced by their deep, well-supported branches. Unlike subspecies A, subspecies B displays a more homogeneous genetic and ecological profile, even though it is further divided into two well-resolved subclades. All strains of subspecies B are related by their MLST allelic profiles (STs 228, 624, 1468, and their respective loci variants), with the majority sharing common isolation sources, such as animal meat or meat processing facilities. From an epidemiological standpoint, the absence of association between subspecies B and human clinical sources, along with the uniform lack of traits commonly found in high-risk lineages (e.g., acquired multidrug resistance and virulence genes linked to treatment failure and increased pathogenicity in enterococcal infections; see reviews (71, 82) for comprehensive overviews) suggest that this subspecies may not pose a significant threat to human health, either as pathogens or as reservoirs of clinically relevant ARGs. However, it is also reasonable to consider that because subspecies B is not commonly found in humans and most molecular epidemiology studies on enterococci are centered around human clinical isolates, our understanding of its niche breadth and ecological roles within a one-health continuum is currently limited.

Drawing from the existing literature and available metadata on strains classified as subspecies B, including the STs represented in our study as well as their single and double-locus variants cataloged in the PubMLST database (https://pubmlst.org/), it appears that this subspecies is predominantly associated with animal-related sources. Notably, it has been isolated not only from meat and meat processing facilities but also directly from cloacal samples of chickens and wild birds such as common kingfishers (*Alcedo atthis*, a species within the Coraciiformes order). These reports are predominantly linked to ST228 (83–85).

The isolation of the ST624 strain from a mouse GIT, as documented in the NCBI under the identifier SAMEA13256522 (DSM111623 biosample), contributes to the understanding that subspecies B may also be associated with non-human mammals, in addition to livestock. Notably, we found no reports of subspecies B-associated STs being isolated from humans, with the sole exception of the ST228 SLV, ST247, found in a subgingival plaque sample from a patient with marginal periodontitis in Norway (86). This observation underscores the rarity of human association for subspecies B, reinforcing the notion of its distinct ecological profile.

The consistent absence of critical genetic determinants associated with *E. faecalis* persistence in hospital environments, enhanced pathogenicity, and treatment failures in HAIs aligns with the universal presence of the CRISPR3-Cas system across all subspecies B genomes examined. Johnson et al. (2021) (5) have reviewed how the evolution of enterococci, particularly *E. faecalis* and *E. faecium*, into MDR hospital-acquired pathogens is largely driven by the accumulation of MGEs, including plasmids, transposable elements, and phages, which serve as vehicles for the spread of ARGs and virulence factors. Given that these traits are predominantly acquired through HGT, and the CRISPR3-Cas systems are known to restrict HGT in *E. faecalis* (87), it seems that subspecies B genomes have been shaped by selective pressures markedly different from those maintaining the high-risk lineages of subspecies A.

An additional facet of this divergence can be seen in the variability of the *ace* gene between subspecies A and B. The Ace protein of *E. faecalis* mediates binding to host extracellular matrix proteins, such as collagen types I and IV and laminin (88, 89), playing a crucial role in the colonization and infection of various tissues (90, 91). Notably, the detection of *ace* genes in subspecies B genomes with significantly lower similarity to the VFDB reference sequence (NP_814829) compared to the *ace* sequences in subspecies A representatives may suggest tropism for different host environments, especially because this gene is known to be highly conserved among *E. faecalis* isolates (92).

Moreover, individual gut bacteria are thought to exhibit divergence patterns aligned with host phylogeny, especially during allopatric speciation, which limits bacterial dispersal between hosts and gradually leads to reproductive isolation (93). Given that enterococci are hypothesized to have been core members of the gut microbiome in the last common ancestor of mammals, birds, reptiles, and insects (94), it is plausible that the divergence between subspecies A and B was driven by host-specific adaptations and reduced gene flow over time. The ubiquitous presence of the CRISPR3-Cas system among subspecies B strains could further support this reduced genetic exchange, offering an alternative or complementary explanation for the reproductive restriction and cladogenesis that culminated in the divergence of *E. faecalis* strains at the subspecies level.

To further investigate the ecological adaptations that differentiate subspecies A and B, within the framework of reverse ecology, we conducted a pangenome analysis to identify genes enriched in each subspecies. This analysis revealed that both subspecies are enriched with genes distributed across the same 20 COG functional categories, suggesting that, in addition to the core genome, each subspecies has developed its own genetic repertoire supporting similar functional capacities. This finding is consistent with their independent evolutionary paths. Notably, subspecies A exhibited a significantly higher proportion of genes involved in carbohydrate transport and metabolism, supporting the hypothesis that subspecies A may be better adapted to environments with greater availability or diversity of carbohydrates, such as the human gut (95–97). In contrast, this suggests that subspecies B may be adapted to more restricted environments where carbohydrate variety is limited, or where alternative carbon sources are more prevalent, such as the guts of animals with more restrictive diets compared to humans, or ecosystems under distinct selective pressures (98). Carbohydrate availability in the host gut has been indicated as a major driver of enterococcal speciation since the emergence of the genus (99), which enhances the plausibility of our findings.

Although these hypotheses align with our primary observations, it is important to note that nearly half of the genes enriched in subspecies B strains were not classified within any COG category. While this finding limits precise inferences regarding subspecies B’s ecological adaptations, it also provides compelling evidence of this lineage’s distinct evolutionary history and adaptive strategies. The significantly higher proportion of unclassified genes in subspecies B, compared to subspecies A, suggests that subspecies B may harbor novel genetic elements that have yet to be fully characterized. These unclassified genes could reflect adaptations to niche-specific pressures, possibly in environments that differ considerably from the well-studied human-associated habitats where subspecies A thrives. Moreover, the absence of functional annotation highlights the possibility that subspecies B is adapted to more specialized or less predictable environments, where distinct selective pressures —such as those found in non-human hosts or natural ecosystems— shape the evolution of unique genetic repertoires. Further research is required to elucidate the roles of these genes and their contributions to subspecies B’s fitness in specific conditions, which will enhance our understanding of *E. faecalis* niche breadth and its complex eco-evolutionary dynamics.

In line with these considerations, it is also relevant to address the observed variability in alleles of MLST housekeeping genes in subspecies B, which may provide additional insights into the ecological divergence between the subspecies. All subspecies B isolates share the same allelic variants of the *gki* and *yqiL* genes, which are key housekeeping loci in the *E. faecalis* MLST scheme (80). Interestingly, Fertner et al. (2011) demonstrated that ST228 — the most frequent ST within subspecies B — had the most phylogenetically distant relationship compared to other *E. faecalis* lineages based on a concatenated phylogeny of the seven MLST housekeeping genes (85). While this study did not directly link the phylogenetic divergence to distinct subspecies, our findings suggest that such genetic distance reflects an ancient and consistent divergence between subspecies A and B, particularly considering that housekeeping genes tend to be highly conserved, with slower rates of evolution compared to other genomic regions (17, 100, 101).

Building on the analysis of genes enriched in each subspecies, we applied a more stringent filter to identify genes present exclusively in one subspecies and absent in the other, across all representative genomes (i.e., clade/subspecies-specific genes, likely synapomorphic traits, and candidate niche-specifying genes) (17, 73). This approach aimed to pinpoint genes potentially associated with the primary niche of each subspecies—those likely acquired by the common ancestor and maintained through periodic selection due to their adaptive value (17, 73). Unlike the broader enrichment analysis, this method focused on genes that may have played a critical role in initiating ecological isolation and, thus, speciation, which is discussed here in light of the concept defined by Shapiro and Polz (2014) as “any stage of the dynamic process of ecological and genetic differentiation” (51).

The prediction of subspecies-specific genes and allelic variants provides valuable targets for understanding the genetic basis of subspecies B’s emancipation and unique ecology. Initially, certain proteins were classified as distinct orthologous groups, thought to be exclusive to each subspecies. However, following reannotation, these proteins were reclassified as products of subspecies-specific allelic variants within the same orthologous group, rather than distinct genes. This finding highlights the potential roles of both gene acquisition or loss events, and sequence variation in pre-existing core genes, in delineating the ecological niches of each subspecies, as supported by Cohan (2002) (102).

Although we cannot confirm whether such allelic variations correspond to actual adaptive mutations or to fixed neutral mutations solely based on these findings, the high degree of non-synonymous substitutions between subspecies A and B sequences suggests potential functional divergence. Even if these variants were neutral in the early stages of ecological speciation, they may have acquired adaptive significance over time, gradually shaped by distinct selective pressures and reflecting the niche specificity of each subspecies (103). It is important to note, however, that not all non-synonymous substitutions necessarily lead to functional changes (104). Therefore, further experimental validation is essential to determine the extent to which these variants contribute to bacterial adaptation (105). Investigating these potential functional changes could significantly enhance our understanding of subspecies-specific ecological adaptation in *E. faecalis*.

Several proteins encoded by subspecies-specific gene variants are potentially linked to stress response and host colonization. One example is the EF1546-encoded protein, which contains a LysM domain, a well-characterized motif in autolysins responsible for binding bacterial cell wall peptidoglycan and facilitating precise catalytic cleavage during daughter cell separation (106). Controlled septum cleavage is critical for defining cell size and chain length in *E. faecalis*, factors directly impacting bacterial fitness (106). Mutations within the LysM domain have previously been associated with elongated cell chains, impaired immune evasion, and reduced virulence in *E. faecalis* (106–109). Given that such phenotypic alterations can profoundly affect bacterial morphology and its interaction with host defenses, the divergence in LysM-containing proteins between subspecies A and B may underlie their differential success in host colonization.

Subspecies-specific sequence divergence was also observed in key transcriptional regulators, which may influence the adaptive responses of *E. faecalis* to varying environmental pressures. The EF1369 gene, encoding a Cro/CI family transcriptional regulator, has been linked to stress tolerance, including survival under high-salt conditions (8% NaCl), acidic environments (pH 2.8), and within mouse peritoneal macrophages (110, 111). Similarly, the EF0876 gene, encoding a Mga helix-turn-helix protein, plays a crucial role in virulence regulation, particularly in host tissue adhesion in response to carbon metabolism (112, 113). Notably, inactivation of EF0876 resulted in a 100-fold reduction in gut colonization in mice in a previous study (113), highlighting its importance for *E. faecalis* fitness in the GIT. Mga family regulators, including key *E. faecalis* regulons like *EbpR* (114), are activated by elevated CO_2_ levels (115, 116), typically found inside mammalian hosts (∼5-6%), as opposed to atmospheric levels in natural settings (∼0.036%) (117, 118). Given the significant role of EF1369 and EF0876 in mediating *E. faecalis* colonization and survival under mammalian host conditions, and considering that these findings are based on subspecies A (i.e., strains derived from the V583 clinical isolate) (110, 113), subspecies-specific sequence divergence in these regulators could impact the fitness of subspecies B under similar selective pressures. This is consistent with the lineage’s unusual isolation from human hosts, and might reflect the genetic basis supporting ecological niche distinctions between the subspecies.

While the adaptive significance of subspecies-specific allelic variants of core genes is less predictable, the acquisition of entirely novel genes encoding new cellular functions is more likely to enable a strain to exploit previously inaccessible resources and establish itself within a new ecological niche (i.e., niche-specifying genes) (17, 51, 102). From this perspective, we highlight our main inferences and speculations regarding the gains and losses of key genetic determinants and the corresponding selective pressures driving the ecological separation of *E. faecalis* subspecies, as discussed below.

Interestingly, the *E. faecalis* V583 EF2063 homologue, identified in the present study as subspecies A-specific, is one of the flanking regions of the CRISPR2 locus in *E. faecalis* genomes (28), whose absence we propose as a genomic marker for subspecies B. The absence of this region in subspecies B suggests that it may have been lost in a common ancestor through a single recombination event, akin to the mechanism of CRISPR1-Cas and CRISPR3-Cas variability described previously (28). While the precise functions of both the EF2063 gene (predicted to encode an AraC-family transcriptional regulator) and the orphan CRISPR2 locus remain unclear, we speculate that their combined loss in subspecies B could reflect a selective pressure favoring the genotype lacking this region, potentially dispensable or deleterious in its primary niche. This hypothesis is supported by the recent description of a wild-type *E. faecalis* strain harboring all three canonical CRISPR loci (CRISPR1-Cas, CRISPR2, and CRISPR3-Cas), leading the authors to conclude that CRISPR genotype variation in the species is likely due to CRISPR loss rather than locus acquisition by HGT (119, 120). However, fitness trade-offs associated with CRISPR variability in *E. faecalis* have been primarily attributed to the presence or absence of functional CRISPR systems, not the orphan CRISPR2 (5). Thus, any further exploration of how CRISPR2 variation contributed to the ecological separation of the subspecies must also consider the role of EF2063 in this process.

Similarly, building on the understanding that functionally related genes are often arranged contiguously in the genome, reflecting potential co-selection under specific environmental pressures (121), we observed that many of the subspecies A-specific genes exhibited synteny, (i.e., based on the gene arrangement of corresponding homologs in the V583 genome sequence) (57). This suggests that they may have been selected together in response to a specific ecological pressure, either gained by the most recent common ancestor of subspecies A or lost in subspecies B.

For instance, the proteins encoded by EF1919 and EF1920 were annotated as a GNAT family acetyltransferase and a C4-dicarboxylate anaerobic carrier DcuC, respectively. While GNAT members are involved in various cellular processes, and this one has not been linked to any specific pathway so far, DcuC carriers are mainly responsible for succinate efflux produced during glucose fermentation (122). Interestingly, most bacteria containing *dcuC* homologs are pathogenic and all known DcuC family members are from enteric bacteria (122). This pattern supports the hypothesis that this locus may be integral to a key carbon and energy metabolic pathway in subspecies A strains, particularly under anaerobic conditions in the gut, the primary habitat of *E. faecalis* where it occurs as both a commensal and opportunistic pathogen (1, 123). In contrast, the absence of these genes in subspecies B could indicate a competitive disadvantage for gut colonization, compared to subspecies A.

Interestingly, many other subspecies A-specific genes seem to encode ecologically relevant determinants involved in *E. faecalis* ability to either colonize, persist or cause infection in a host environment. The EF1672 gene, encoding an ABC superfamily protein permease, was also exclusively and universally detected within subspecies A genomes. Notably, transcriptomic data have supported this gene’s crucial involvement in *E. faecalis* physiological adaptation during the course of infection in a mammalian host model, being hypothesized to be one of the core genes required for the species to thrive within a host, and thus a potential target for antimicrobial agents and vaccines to treat and prevent enterococcal infections (124).

Another example implies that subspecies B strains might lack or have a reduced ability to utilize sucrose as a carbon and energy source. This is supported by the exclusive detection of determinants of two operons encoding a sucrose PTS (phosphoenolpyruvate: sugar phosphotransferase system) transporter (EF1602) and sucrose metabolism (EF1603, also named *scrB-1*) within subspecies A strains. Previous reports have shown that both loci likely contribute to virulence since they were up-regulated in *E. faecalis* strains grown in human urine (125), where sucrose levels can be increased in high-sugar diets (126). Knock-out mutants of EF1603-04 showed reduced virulence in a *Caenorhabditis elegans* infection model (127), which corroborates the differential role of these loci in *E. faecalis* pathogenesis associated with sucrose’s contribution to growth in infection. Additionally, studies have demonstrated that sucrose utilization enhances the expression of *E. faecalis* virulence-associated determinants and the production of biofilm matrix components (e.g., eDNA and EPS) in biofilms (128). Altogether, these findings support the genetic basis of subspecies A pathogenic potential in contrast to subspecies B by highlighting the contribution of sucrose uptake and metabolism in infections caused by *E. faecalis*, which seems consistent with this species (specifically, subspecies A) being a notorious agent of UTIs and aligned with the relevance of biofilm formation in the context of HAIs (82).

Additionally, the subspecies A-specific EF1138 gene encodes an oxidoreductase of the aldo/keto reductase family, strongly related to *E. faecalis* V583 stress response induced by bovine bile exposure, being potentially implicated in bile salt modification (129, 130). Bile tolerance mechanisms are critical for bacterial survival and colonization of the GIT, where bile’s emulsifying action poses a challenge (131). The absence of EF1138 in subspecies B may suggest that bile is not a strong selective pressure in its core ecological niche, or that subspecies B has evolved alternative mechanisms to tolerate bile exposure or to cope with different bile salt compositions, which includes the possibility of adaptation to a different host range (132–134).

Considering that most studies on *E. faecalis*-host interactions have focused on its pathogenic potential in human infections and often rely on conventional animal models, primarily mammals (e.g., mice and rabbits), it is important to acknowledge that conclusions from these studies are inherently biased by the unique physiology of each host species (135). While the absence of subspecies A-specific genes in subspecies B suggests reduced fitness in humans and other mammalian hosts, it remains uncertain whether these genes are essential for colonization and survival across a broader host range, particularly in non-mammalian vertebrates (97, 136). This knowledge gap limits our understanding of subspecies B’s ecological niche and adaptability.

The isolation sources of subspecies B, which include wild birds, poultry, and meat processing environments, suggest a potential adaptation to hosts or environments beyond humans (83–85). Although the specific ecological role of birds in the lifecycle of subspecies B remains uncertain, previous studies support the hypothesis that wild migratory birds act as reservoirs for genetically diverse *E. faecalis* strains. Some of these strains are prevalent in poultry, indicating possible transmission routes from wild to domestic birds (16, 137). These findings reinforce the idea that subspecies B may have evolved to thrive in non-mammalian hosts, particularly avian species, whose gut microbiota composition is consistently distinct from that of mammals (138).

Furthermore, according to conclusions of a study by Chaillou and colleagues on meat microbiota (139), subspecies B’s common isolation from meat products and processing environments likely points to two primary sources of contamination: (1) animal-derived microbiota, particularly from the skin or gut, which may include *Enterococcus*, and (2) environmental psychrotrophic bacteria, largely represented by *Firmicutes* associated with water reservoirs, plants, or plant-derived animal feed (139). These hypotheses are not mutually exclusive in the context of *E. faecalis*, but rather suggest multiple potential dissemination pathways. This also implies that subspecies B may have undergone selective adaptations to thrive in such environments, where particular pressures could have driven and maintained its ecological and genetic differentiation over time (51).

Our efforts to obtain further insights into the unique ecological niche of subspecies B focused on the functional annotation of its specific genes, a reverse ecology approach (17, 73). While many of these genes encode proteins with uncharacterized functions, functional predictions were made based on putative homologs and conserved domains. Despite the significant homology between subspecies B sequences and their closest matches in the UniProtKB database, strongly supported by low E-values in most cases (54), notable differences in sequence identity were observed, making functional predictions less definitive. Nevertheless, the detection of conserved protein signatures provided valuable insights into potential ecological roles. Interestingly, most of the predicted homologs were derived from taxa commonly associated with meat microbiota and the contamination routes proposed by Chaillou et al., such as *Vagococcus*, *Carnobacterium*, *Leuconostoc*, various enterococcal species, and others (139). This correlation suggests that these genetic determinants may confer adaptations to shared ecological niches, potentially aiding subspecies B in coping with the selective pressures of these environments.

One of our most compelling findings was the detection of sequences potentially encoding the toxin-antitoxin (TA) protein components HicA and HicB, along with a ClpP/crotonase-like domain-containing protein. These elements may play a coordinated role in stress responses linked to environmental conditions such as nutrient deprivation and heat shock (140, 141). It has also been suggested that TA systems, including HicAB, are typically abundant in free-living bacteria but tend to be lost in host-associated prokaryotes (142, 143). This raises the possibility that subspecies B’s recurrent isolation from meat products could be influenced by environmental contamination routes, potentially involving water, plants, or soil.

Additionally, the ubiquitous detection of the CRISPR3-cas system within subspecies B further supports the contribution of environmental pressures, as functional CRISPR-Cas systems are known to provide defense against lytic phages, which are particularly abundant in water and soil environments (144–146). These environments could impose strong selection, contributing to the maintenance of CRISPR-mediated defense mechanisms. Although the CRISPR3-cas system was not exclusive to subspecies B, we observed that its Cas proteins (Cas1, Cas2, and Csn2) clustered distinctly from those of subspecies A, showing greater similarity to those found in *Vagococcus elongatus*, a species originally isolated from a swine-manure storage pit (147), which aligns with the transmission routes involving either animal or environmental reservoirs to meat microbiota, as proposed by Chaillou and colleagues (139). This divergence could reflect adaptations to different ecological pressures, though further investigation is needed to clarify the specific ecological role of these systems.

We also identified subspecies B-specific traits that may be involved in the uptake and metabolism of alternative carbohydrates when glucose is scarce, which could suggest an adaptive advantage in diverse environments. Notably, we detected protein clusters resembling the IIABC component (BglF-like permease) of a PTS beta-glucoside-specific system and a PRD (PTS regulation domains)-containing protein (LicT-like transcriptional antiterminator) (148, 149). These elements likely participate in the transport and positive regulation of genes linked to beta-glucoside metabolism, responding to substrates like salicin and arbutin found primarily in plant material but rarely in mammals (150).

Moreover, we observed the exclusive presence of putative carbohydrate-binding proteins in subspecies B, such as the GlcNAc-binding protein A (GbpA) and a WxL domain-containing protein. The GbpA homolog contains domains frequently associated with cellulose- and chitin-binding capabilities, polysaccharides predominantly found in plants and arthropods, or fungi, respectively (151, 152). In *Vibrio cholerae*, GbpA facilitates adhesion to zooplankton chitinous surfaces in aquatic habitats and to GlcNAc moieties on human intestinal epithelium (153). This dual functionality strongly suggests an exaptive process where a protein initially evolved for environmental persistence (e.g., attachment to chitin) later adapted to facilitate host colonization (154). By analogy, a similar exaptation in *E. faecalis* subspecies B could enable these bacteria to persist in environmental reservoirs (potentially associating with arthropods or adhering to plant cell walls) and also to colonize animal gut surfaces via GlcNAc binding, supporting its ecological adaptability across diverse environments (153, 154).

The WxL domain-containing proteins, commonly associated with adhesion to cellulose and xylan in other Gram-positive bacteria, such as *Enterococcus faecium* (155), reinforce this notion. These polysaccharides, abundant in plant cell walls, are often present in animal diets as dietary fibers. Interestingly, the ability of gut bacteria to adhere to these fibers confers a competitive advantage, especially in environments where plant-based feeds are supplemented with xylanase, facilitating the growth of beneficial microbes like lactobacilli (155, 156). Thus, the exclusive presence of a WxL domain-containing protein in subspecies B suggests that these bacteria might have adapted to adhere to dietary fibers in the gastrointestinal tract of animals, potentially contributing to their isolation from meat products.

Collectively, the subspecies B-specific GlcNAc-binding and WxL domain-containing proteins exemplify how exaptation might facilitate bacterial adaptation to shifting habitats, supporting the two potential transmission routes proposed by Chaillou et al. (2015). These traits could either enable direct environmental persistence, leading to contamination of meat products, or facilitate colonization of animal hosts, with subsequent transmission through the food production chain. However, it is important to note that identifying these proteins as exaptive traits remains speculative and requires further investigation for confirmation. Nevertheless, considering such possibilities underscores the adaptive potential of subspecies B and highlights the value of genomic surveillance across diverse settings to understand the evolutionary mechanisms driving *E. faecalis* adaptation, especially from a One Health perspective.

Our findings also revealed specific subspecies B traits that suggest the existence of an unique repertoire of surface proteins, including several containing the LPxTG cell wall anchor motif, a feature typically associated with cell surface adhesion in Gram-positive bacteria (157). This repertoire appears to include proteins that may mediate adhesion to various substrates, as inferred from other conserved domains identified in their sequences. For example, one protein exhibits the fimbrial isopeptide formation D2 domain, which may facilitate pili subunit cross-linking and mediate surface adhesion to host structures (158, 159). Another predicted protein contains a T-Q ester bond domain, an unusual bond type thought to stabilize pilin subunits and commonly found in cell surface adhesion proteins of Gram-positive bacteria (160), similar to a known collagen adhesin from *Bacillus cereus*.

Moreover, we identified three proteins with MucBP domains, matching putative adhesins encoded by other enterococcal species. Notably, two of these proteins showed the highest similarity to a sequence from *Enterococcus phoeniculicola* ATCC BAA-412, a strain isolated from the uropygial gland of wild Red-billed Woodhoopoes (*Phoeniculus purpureus*) (161). This finding supports our earlier suggestion that subspecies B may be adapted to avian hosts, as the presence of proteins homologous to those in a bird-associated *E. phoeniculicola* highlights the potential role of birds as hosts or reservoirs for subspecies B. MucBP domain-containing proteins are generally linked to adherence in the mucosal environment of the GIT, providing a potential mechanism for subspecies B to colonize this niche. Notably, previous studies have identified these proteins as common in other commensal and environmental Gram-positive species, such as *Lactobacillus plantarum* (162–164). This suggests that subspecies B might exhibit similar ecological flexibility, capable of thriving in a range of environments, including the GIT of animals.

Finally, two subspecies B-specific protein clusters were similar to predicted ABI family (Rce-like) proteins, both classified as type II CAAX prenyl endopeptidases. ABI proteins in enterococci have been previously associated with defense against phage infections (165), but there is also evidence suggesting their involvement in bacteriocin self-immunity mechanisms (166). This suggests that subspecies B might be able to produce additional bacteriocins. Unlike antibiotics, bacteriocins exhibit a narrow antimicrobial spectrum, targeting closely related bacterial strains (167). Such specificity implies that bacteriocin production might confer a competitive advantage by selectively inhibiting competitors within the same ecological niche. Additionally, bacteriocin production in Gram-positive bacteria is often linked to the transition from the logarithmic to stationary growth phase (168), highlighting their role as potent killing agents in resource-limited environments. Thus, the presence of subspecies B-specific proteins potentially involved in bacteriocin self-immunity hints at the possibility that this clade may possess a distinct competitive advantage in microbial communities, either in host-associated microbiomes or in environmental settings (169, 170).

Our findings suggest that the niche breadth of *E. faecalis* is likely underestimated, potentially encompassing a range of unexplored or insufficiently studied hosts and environments, which may explain the infrequent reporting of subspecies B-associated STs. By employing a reverse ecology approach, we have provided several insights into the hypothetical ecological properties and potential niches of subspecies B, aligning with current guidelines for the taxonomic description of prokaryotes from genome data (78).

Nonetheless, the limitations inherent to our methods must be acknowledged. Specifically, the focus on clade-specific genes to identify past adaptations conserved through periodic selection inevitably overlooks other genetic determinants that may have contributed to the lineage’s evolution and differentiation, as highlighted by Lassalle et al. (2015). Additionally, experimental validation is essential to confirm the speculated roles of clade-specific genes or alleles and corresponding selective pressures in driving the ecological differentiation between *E. faecalis* subspecies A and B, following similar approaches used in related studies (73, 99, 171, 172).

Despite these limitations, our study highlights how genome-based exploratory approaches, utilizing publicly available data, can provide novel insights into the eco-evolutionary dynamics of bacterial species while simultaneously revealing important knowledge gaps. In the present study, analyzing CRISPR system genetic patterns across diverse genomes (spanning a broad range of STs, isolation sources, etc.) within the operational definition of a single species enabled us to identify and partially characterize a distinct cluster of *E. faecalis* strains, corresponding to a phylogenetically, ecologically, and taxonomically cohesive unit —subspecies B, using approaches proposed by previous studies (17, 51, 73, 78).

Although previous works have depicted the phylogenetic prominence of subspecies B members, particularly ST228 strains, compared to other *E. faecalis* clusters (83, 85), neither the subspecies-level classification of this lineage, nor its positioning within the broader landscape of *E. faecalis*’s population structure have been acknowledged. Addressing this gap, our findings reinforce the need to reassess *E. faecalis*’s population structure in light of hidden genetic diversity in under-sampled environments and hosts (e.g., nonhuman isolates), as suggested by other studies (16, 123). Given the role of generalist species in microbial dispersal and their propensity to give rise to specialists confined to a narrower range of habitats (173, 174), it is plausible that the current view of *E. faecalis* as a species with little phylogenetic divergence and lacking a multiclade structure (15, 175) stems from an underrepresentation of its true biological diversity. This underrepresentation likely obscures the existence of host- or environment-specific clades in advanced stages of speciation, such as subspecies B.

Expanding intraspecific genomic surveillance to include isolates from nonhuman hosts and nonclinical environments will foster a more realistic understanding of the species’ population structure. Such efforts are critical from a One Health perspective, especially given previous evidence that *E. faecalis*’s adaptation to the hospital environment may be a byproduct of its evolution in a wider range of ecological niches (6, 99, 175). Investigating the existence of niche-specific *E. faecalis* clades can help predict and monitor potential adaptive novelties that may precede the emergence of high-risk lineages, allowing for more targeted prevention and control strategies aimed at curbing the spread of these genetic determinants and associated lineages (176, 177).

While opportunities for genetic exchange between subspecies A and B are likely provided in overlapping compartments of their multidimensional niche breadths (178), it remains to be elucidated whether and how gene flow between individuals of these clades is maintained.

## CONCLUSIONS

In this study, we unexpectedly identified a genetically related cluster of *E. faecalis* strains that lack the CRISPR2 locus, previously considered universally present in the species. Remarkably, this cluster represents a distinct subspecies, which we designate as subspecies B - an unprecedented finding, as no subspecies supported by formal genome-based taxonomic criteria has been previously described in *E. faecalis*. Our pangenome and reverse ecology analyses suggest that differences in carbohydrate availability in the host gut are likely key selective pressures driving the ecological separation between subspecies A and B. While subspecies B appears to have limited fitness in the mammalian gut and lacks genetic determinants associated with HAIs and multidrug resistance, it may have evolved under distinct host physiologies, potentially adapting to avian species. Furthermore, subspecies B harbors a unique genetic repertoire that likely enables its persistence in natural environments, such as water reservoirs, plants, or plant-derived animal feed, which may explain its frequent isolation from meat products and processing facilities.

These findings challenge the previously established conception of the population structure of *E. faecalis*, revealing hidden phylogenetic diversity present in less-studied environments and hosts. From a One Health perspective, exploring intraspecific genomic heterogeneity, including nonhuman isolates, is essential for unraveling the genetic basis and selective pressures that drive the emergence of bacterial variants. Understanding whether these variants pose imminent public health threats, as well as identifying potential dissemination routes and environmental and host reservoirs, is crucial for developing targeted strategies to prevent and control the spread of emerging resistant and virulent genotypes.

## ACKNOWLEDGMENTS

This work was supported in part by Conselho Nacional de Desenvolvimento Científico e Tecnológico (CNPq: Projects PQ/CNPq 312801/2020-3 and RAM/CNPq 408725/2022-2), Instituto Nacional de Pesquisa em Resistência Antimicrobiana (INPRA Project: INCT/CNPq 465718/2014-0); Fundação de Amparo à Pesquisa do Estado do Rio de Janeiro (FAPERJ: Projects E-26/211.554/2019, E-26/201.084/2021 and E-26/210.064/2020), and Coordenação de Aperfeiçoamento de Pessoal de Nível Superior (CAPES)–Finance Code 001

